# Bacteriophages encoding human immune evasion factors adapt to livestock-associated MRSA through rounds of integration and excision

**DOI:** 10.1101/2021.04.29.441770

**Authors:** Helena Leinweber, Raphael Sieber, Jesper Larsen, Marc Stegger, Hanne Ingmer

**Affiliations:** Department of Veterinary and Animal Sciences, University of Copenhagen, Stigbøjlen 4, 1870 Copenhagen, Denmark; Department of Bacteria, Parasites and Fungi, Statens Serum Institut, Artillerivej 5, 2300 Copenhagen, Denmark

**Keywords:** Livestock MRSA, CC398, Sa3Int, phi13, Φ13, excision, alternative integration, tyrosine recombinase, attP, attB

## Abstract

In recent years there has been an increase in human infections with methicillin-resistant *Staphylococcus aureus* (MRSA) originating from livestock and strains carrying bacterial viruses of the Sa3int-family have disseminated into the community. Sa3int phages express immune evasion factors and are common in human staphylococcal strains. As the bacterial attachment site (*attB*) for Sa3int phages is mutated in livestock-associated strains, the integration frequency is low and a key question is how the phages are established. Here we show that Sa3int phages adapt to alternative bacterial integration sites by mutating the phage attachment sequence, *attP*, leading to enhanced integration at these sites. Using a model strain carrying the mutated *attB_LA_* of livestock-associated strains we find that once established, the Sa3int phage, Φ13 is inducible with release of heterogenous phage populations carrying mutations in *attP* that in part increase homology to alternative integration sites or *attB_LA_*. Compared to the original phage, the adaptive mutations increase phage integration in new rounds of infection. Also, Sa3int phages induced from livestock-associated outbreak strains reveal mutated *attP* sequences. We suspect that promiscuity of the phage-encoded recombinase allows this adaptation and propose it may explain how phages mediate “host jumps” that are regularly observed for staphylococcal lineages.

## Introduction

*Staphylococcus aureus* colonizes both humans and animals and its preference is associated with the content of mobile genetic elements ^1^. One example is the prophages of the Sa3int family. These bacterial viruses are found in most human strains of *S. aureus* where they express one or more immune evasion factors believed to facilitate human colonization as well as to promote human-to-human transmission ^2,3^. In contrast, the methicillin-resistant *S. aureus* found in livestock (LA-MRSA) commonly lack Sa3int phages ^4,5^. In fact, LA-MRSA of the CC398 lineage have been derived from human-associated strains which, subsequent to a jump from humans to animals, lost the Sa3int prophage ^4^.

Despite host preference, there is a growing number of human infections with LA-MRSA and in 2019, they accounted for 32% of all new MRSA cases in Denmark (DANMAP, 2019). People with occupational livestock contact are most at risk ^6–8^ and the infections appear equally severe as those caused by human-associated strains ^9^. Although human infections with LA-MRSA are considered to be the result of spillovers from livestock, there have been examples of transmissions between household members as well as into community and healthcare settings ^3,7,8^. Importantly, such transfer events were associated with LA-MRSA strains carrying prophages of the Sa3int family ^3,7,8,10^. Since 95% of tested Danish pig herds are positive for LA-MRSA (DANMAP, 2019), establishment of Sa3int phages in these strains may pose an increased risk of community spread of LA-MRSA strains.

Integration of Sa3int phages in *S. aureus* occurs through orientation-specific recombination between identical 14bp phage and bacterial core attachment sequences (*attP* and *attB*, respectively) and is mediated by the phage-encoded tyrosine recombinase, Int ^11,12^. In livestock strains, the sequence corresponding to *attB* has two nucleotide changes as underlined 5’-TGTATCC<u>**G**</u>AA<u>**T**</u>TGG-3’ (*attB_LA_*). These substitutions do not alter the amino acid sequence of the β- hemolysin encoded by *hlb* in which *attB* is located, but significantly decrease the ability of Sa3int phages to insert at this location ^13^. Accordingly, in LA-MRSA strains Sa3int prophages are mostly located at variable positions in the bacterial genome but occasionally also in *attB_LA_* ^7,13–16^.

The ability of *S. aureus* to change its preference for human or animal hosts has been observed several times. Such “host jumps” are thought to arise from “spillover” events where infections of less preferred hosts are followed by host adaptation ultimately leading to colonization ^7,17^. Host adaptation often involves acquisition or loss of mobile genetic elements such as prophages ^1^ but little is known of the molecular events involved. Using massive parallel sequencing we have examined Sa3int phages excised from alternative integration sites and find phage populations with variable *attP* sequences of which a greater part increase resemblance to the bacterial attachment sequence. Infections of naïve strains carrying the *attB_LA_* site with such phage pools result in increased phage integration. Our results explain how Sa3int phages, by adapting to alternative integration sites in LA-MRSA strains, can establish in these strains that ultimately may be more successful at colonizing and infecting humans and to disseminate in the human population.

## Results

### Sa3int phages are adapting to alternative *attB* sites of LA-MRSA CC398

In a recent study, 20 LA-MRSA CC398 strains from pigs and humans in Denmark were isolated and found to contain Sa3int prophages. In these strains the prophages were located at one of five different genomic locations (variant I-VI) ^7^ and the 14bp bacterial integration site carried two nucleotide mismatches (designated *attB_LA_*) compared to the one found in human strains in other studies of LA-MRSA strains ^13,15,16^. In the LA-MRSA CC398 genomes we determined the sequences flanking the prophage (*attL* and *attR*) and through comparisons with strains that lack the prophage, we deduced the corresponding *attB* sequences (Table 1). In all cases except one (variant V), the *attL* sequences differed from *attR*. This indicates non-matching *attB* and *attP* sites, as otherwise *attR* and *attL* would be identical, as seen with the original *attB*-site in *hlb* of *S. aureus* 8325-4.

**Table 1.**
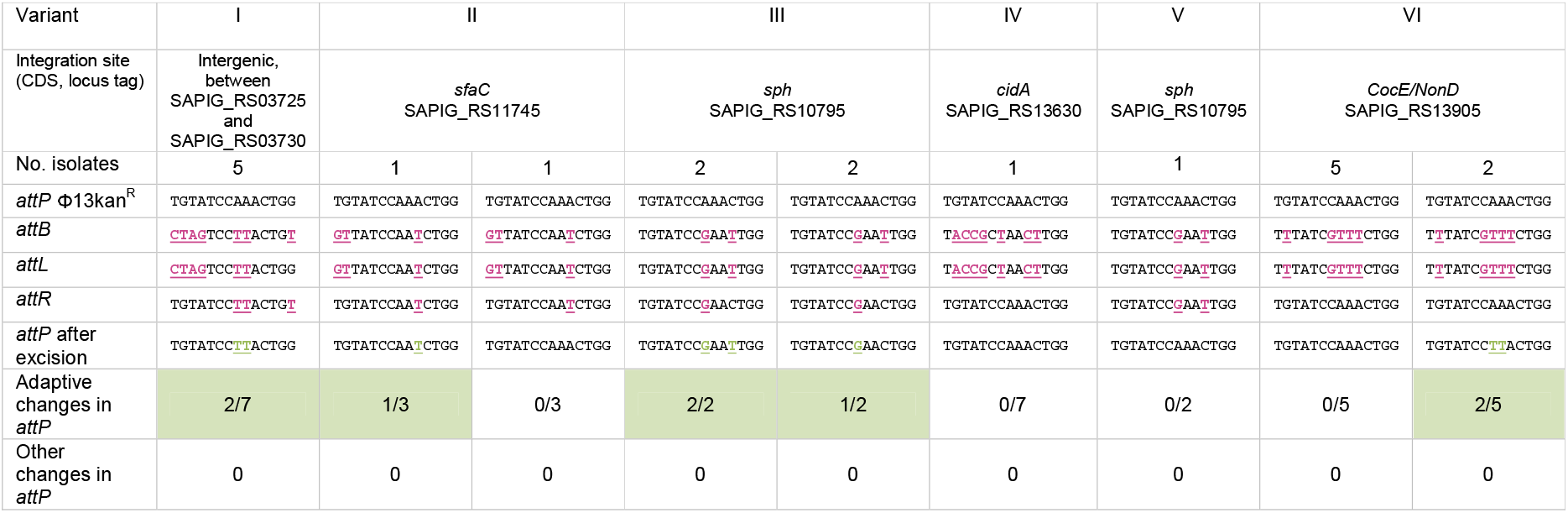
Comparison of *attP*, *attB*, *attL*, and *attR* sites of Sa3int phages from 20 LA-MRSA CC398 isolates. Magenta underlined nucleotides indicate mismatches between the Φ13kan^R^ *attP* core sequence (5’-TGTATCCAAACTGG-3’) and the *attB* site for each LA-MRSA CC398 isolate. Underlined nucleotides in green indicate putative adaptive changes in the *attP* site that mimic the *attB* site under the assumption that the phages contained the *attP* site core sequence upon integration into the different LA-MRSA CC398 genomes. Integration sites refer to annotated genes in *S. aureus* ST398 reference strain S0385 (GenBank accession no. NC_017333).

To examine if mismatches between *attL* and *attR* affected excision of the prophage, we induced the lysogens with mitomycin and observed that in all strains the phages could be excised. From the resulting phages we determined the *attP* sequences using PCR amplification and Sanger sequencing (Table 1). For eight phages (one isolate of variant II, variant IV, variant V and five isolates of variant VI), the *attP* sequences were identical to that of the model Sa3int phage Φ13 ^11^, showing that in these cases integration in the variant *attB* sites did not affect the *attP* sequence of the excised phage. In the remaining 12 phages however, mutations had arisen in the phage *attP* sequences. Importantly, in all cases the changes increased the sequence similarity between *attP* and the alternative *attB* site of the livestock-associated strains, as indicated in Table 1. These results indicate that Sa3int phages may be promiscuous with respect to both integration and excision and that integration of prophages at alternative bacterial attachment sites may alter the phage in such a way that its *attP* sequence bares greater resemblance to alternative *attB* sequences.

### Phage integration at multiple locations in a model strain carrying *attB_LA_*

To examine the interactions between Sa3int phages and LA-MRSA strains in greater detail, we employed a derivative of *S. aureus* NCTC8325-4, designated *S. aureus* 8325-4attBmut, which contains 2bp point-mutations in *hlb* to create the *attB_LA_* of the LA-MRSA CC398 lineage ^13^. With this strain we performed liquid infection with Φ13kan^R^, a derivative of the Sa3int phage, Φ13 that encodes the staphylokinase (*sak*) but in which the immune evasion virulence genes *scn* and *chp* are replaced by the kanamycin resistance cassette *aph*A3 ^13^.

From eight independent lysogenization experiments we selected 22 lysogens being resistant to kanamycin. Alternative integration sites were confirmed for 20 of the lysogens by PCR (*hlb*+, *sak*+) and two lysogens harbored the phage in the mutated *hlb* site (*hlb*−, *sak*+) (Supplementary Figure S1). The 22 isolates were whole-genome sequenced and analysis revealed 17 different integration sites for Φ13kan^R^ in *S. aureus* 8325-4attBmut that were widely distributed across the bacterial chromosome (Supplementary Figure S2) and with the *attB* sequences listed in Figure 1. The integrations occurred in both non-coding and coding regions and were independent of transcriptional orientation.

**Figure 1.**
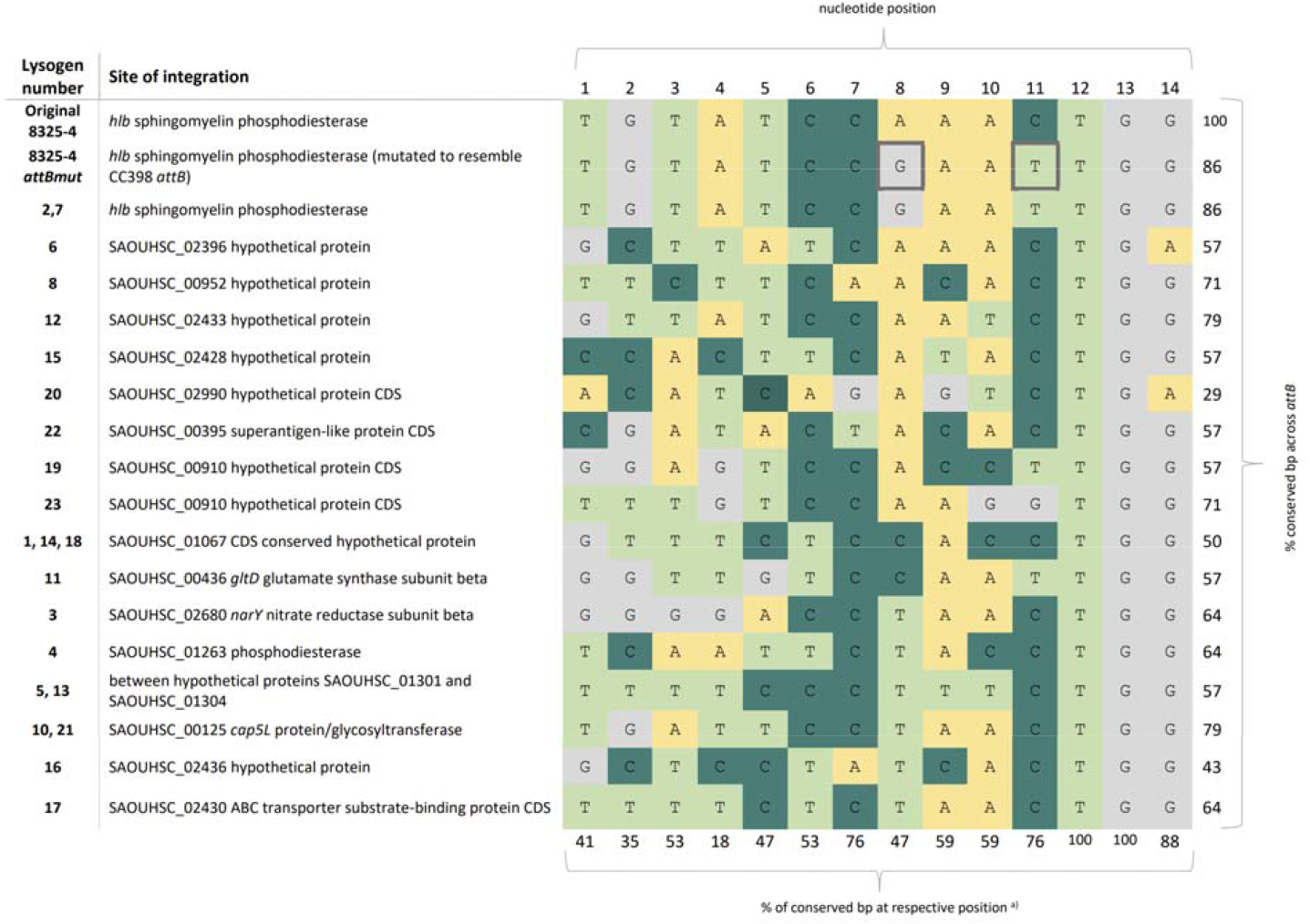
Alternative integration sites of Φ13kan^R^ in *S. aureus* 8325-4attBmut. The core *attB* sites are presented by color-coding of the different base pairs (A = yellow, C = dark green, T = light green, G = grey). The mutated base pairs in *hlb* representing *attBLA*, in the recipient strain are indicated by a bold frame. ^a)^ The percentages in the bottom row correspond to the proportions of conserved nucleotides in the 17 alternative *attB* sites found in the 22 lysogens with respect to the original *attB* in 8325-4.

When comparing the 14bp sequences of all alternative *attB* sites (Figure 1), they showed 29-86% homology compared to the original *attB* core sequence in the *hlb* gene. However, the last three base pairs (5’-TGG-’3) were highly conserved, being present in 20 out of 22 *attB* sites with lysogens 6 and 20 being the exceptions. The nucleotides G at position 8 and T at position 11 signifying the *attB_LA_* compared to *attB*, were not found in the same combination in any of the 17 *attB* sequences. Based on the conserved base pairs between the alternative *attB*-sites, we searched the chromosome of *S. aureus* NCTC8325 for the presence of 5’-NNNNNNCWNNCTGG-’3 (where W = A/T) and obtained more than 700 hits. Thus, there appears to be a multitude of potential integration sites in the staphylococcal genome.

Three of the alternative *attB* locations were observed as integration sites in lysogens obtained in independent lysogenization rounds, i.e. SAOUHSC_01067 CDS conserved hypothetical protein (lysogens 1,14 and 18), the intergenic region between open reading frames encoding hypothetical proteins SAOUHSC_01301 and SAOUHSC_01304 (lysogens 5 and 13) and SAOUHSC_00125 *cap5L* protein/glycosyltransferase (lysogens 10 and 21). As clonality can be excluded, these integration events show that there is some preference in selection of integration site when the bona fide *attB* sequence is mutated. However, when we screened the 300bp flanking regions of the alternative *attB* sites we found no common patterns in terms of sequence composition or distance of inverted repeats relative to the alternative *attB* core sequences (Supplementary Figure S3 and S4). Thus, it is still unclear why some integration sites are preferred over others.

### Phage evolution following excision from alternative integration sites

Similar to what we had observed for Sa3int phages in livestock-associated strains, we found that mitomycin C induced Φ13kan^R^ from all lysogens established in the 8325-4attBmut strain with the number of phage particles varying between 5×10^3^ plaque-forming units (pfu)/ml and 4×10^6^ pfu/ml (Supplementary Figure S5). This represents up to 1000-fold decrease in induction efficacy compared to the 6×10^6^ pfu/ml obtained when the phage was induced from its integration site in the non-mutated *attB* of *S. aureus* 8325-4 (8325-4phi13kan^R^ control). Spontaneous phage release was also detected for many of the lysogens ranging from 2×10^1^ to 3×10^3^ pfu/ml compared to 1,0×10^4^ pfu/ml for the 8325-4phi13kan^R^ control (Supplementary Figure S5).

To examine the integration and excision process of Φ13kan^R^ at the alternative integration sites, we determined the *attL* and *attR* from the genome sequences of the lysogens and deduced the alternative *attB* sites by comparing with sequences prior to integration of the phage. In addition, we determined the *attP* sequences by induction of the lysogens and amplicon sequencing of PCR products obtained on phage lysate with primers spanning *attP* (sequencing depth range 10.000-180.000, average 100.000).

For the majority of the lysogens (Table 2, part A), *attL* was identical to *attB*, and *attR* was identical to *attP* as can be observed by the bold red nucleotides marking the nucleotide differences in the alternative *attB* site sequences compared to the original *attB*. For these lysogens, the integration cross-over likely occurred at the 5’-TGG-3’ (Supplementary Figure S6a). For the remaining lysogens (Table 2, part B), both *attL* and *attR* displayed sequences matching the alternative *attB* site with *attL* matching the 5’-end and *attR* the 3’-end. In these cases, the integration cross-over events may have occurred at variable positions within the core sequences (Supplementary Figure S6b).

**Table 2a and b.**
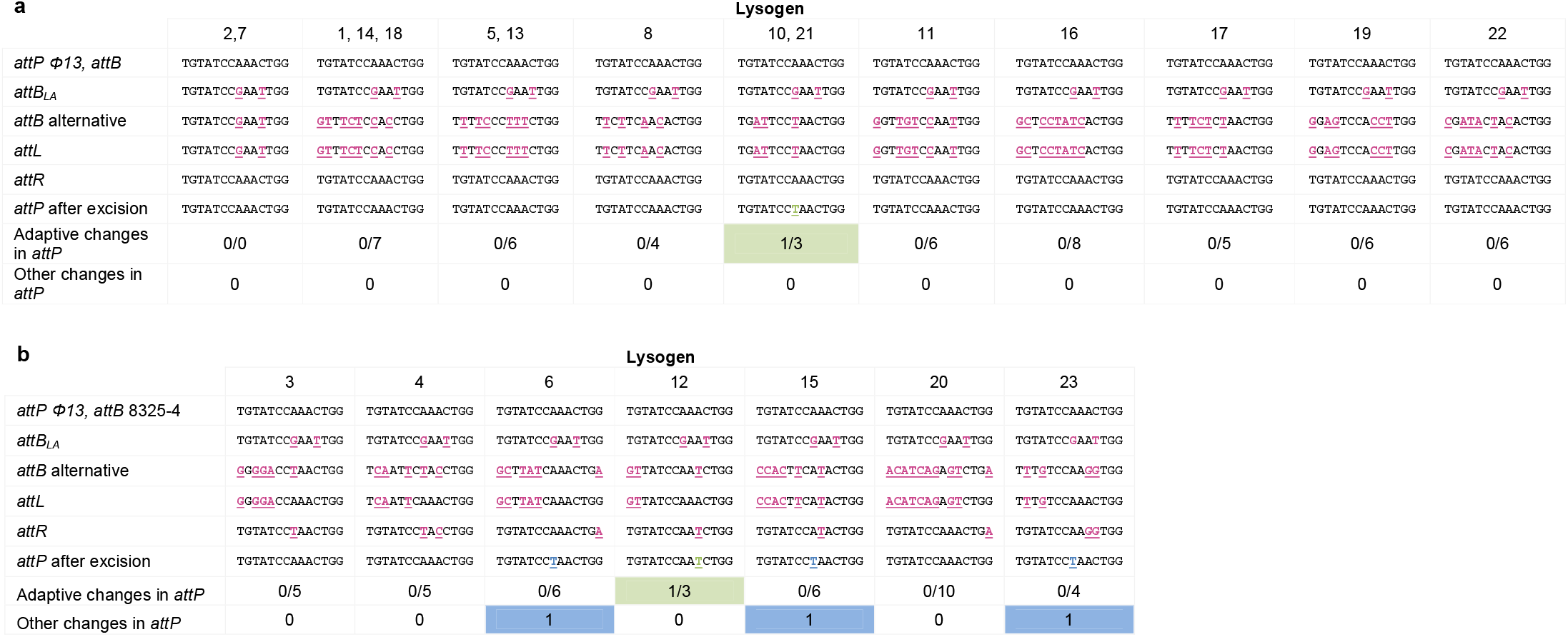
Comparison of *attP, attB, attL*, and *attR* sites of Φ13kan^R^ in 8325-4attBmut. Magenta underlined nucleotides indicate mismatches between the *attB* site in 8325-4 and the *attB* site for each lysogen. Green underlined nucleotides indicate adaptive changes in the *attP* site that mimic the *attB* site. Blue underlined nucleotides indicate other changes in the *attP* site. The threshold for variant calling was set to 50%. Part A includes the lysogens, where *attL* matches *attB* and *attR* matches *attP*. Part B includes the lysogens where parts of *attL* and *attR* both match *attB* and *attP*.

When assessing *attP* by amplicon sequencing we observed remarkable sequence variation at single nucleotide positions in more than 40% of the phage populations obtained from 9 of the lysogens (Figure 2). When comparing these changes to the sequence of the bacterial integration site from which the phage was derived, we saw that in five instances (lysogens 3, 10, 12, 17 and 21) the excised phages displayed adaptation to the alternative *attB* site by adopting a nucleotide of the alternative *attB* sequence (Figure 2). Phages from lysogens 6, 7, 15, and 23 also displayed single nucleotide substitutions in *attP* but without matching the alternative *attB* sequences. These may result from mismatch repair during excision.

**Figure 2.**
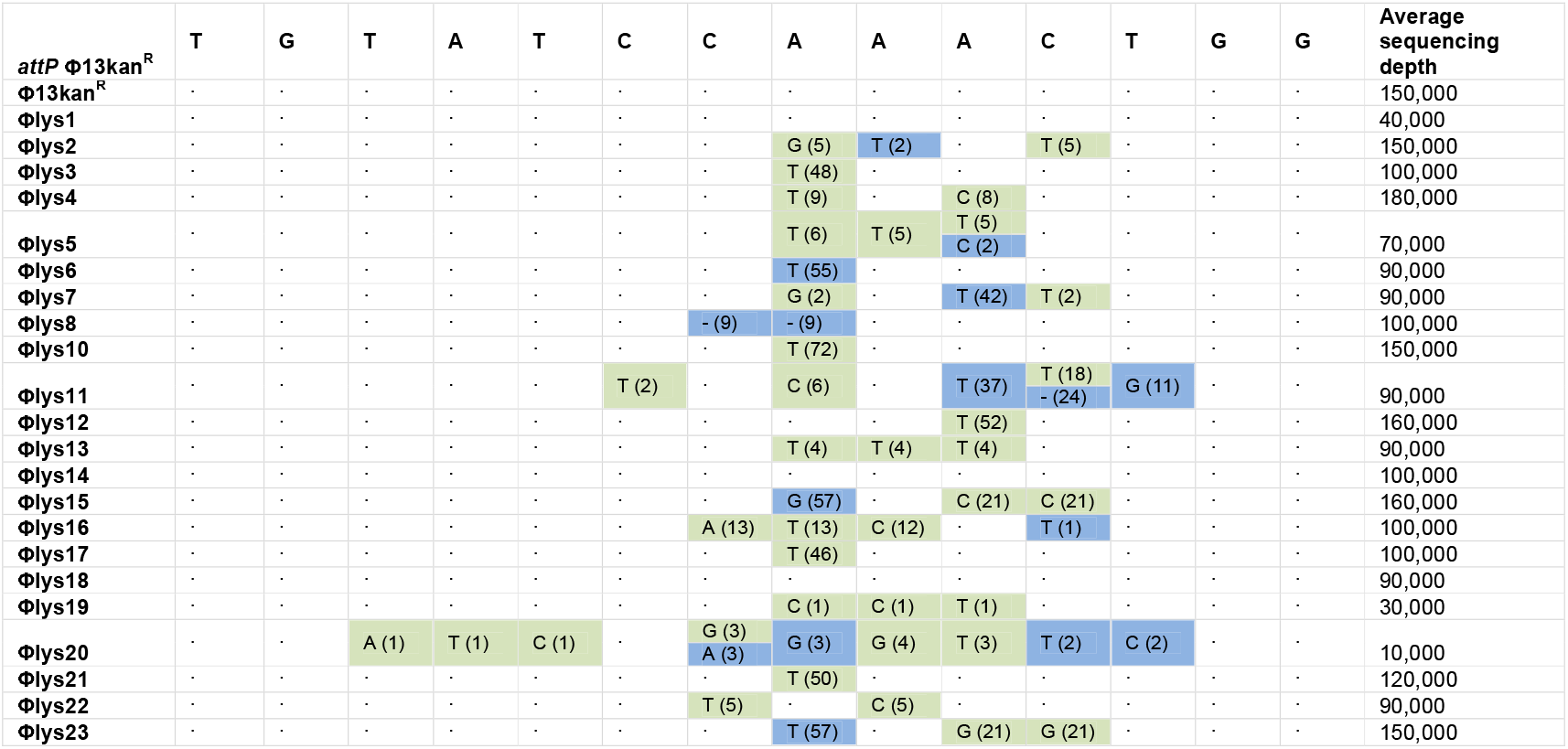
Variant nucleotides and respective frequency (%) of the *attP* sequences after excision of the phage as determined by amplicon sequencing. The green shading of the percentages indicates adaptive changes in the *attP* site that mimic the respective alternative *attB* site. Blue shading of the percentages indicates other changes at this position. A hyphen (-) indicates that no nucleotide was detected at this respective position. Dots indicate conservation of the base pair compared to *attP* in Φ13kan^R^, the sequence of which is indicated in the first row. Note that for Φ13kan^R^, Φlys1, Φlys14 and Φlys18 no variants with frequencies >1% were detected across the entire *attP* sequence.

The adaptability of the phage to the alternative integration sites was even more pronounced when all sequence variation >1% was scored (Figure 2). Importantly most of the excised phage pools contained variants with sequence changes adopting the nucleotides of the alternative *attB* sequences and multiple sequence variations occurred within the individual pools (Figure 2, green). Notable exceptions were lysogens 1,14 and 18, where no variants >1% were observed. In these lysogens, Φ13kan^R^ had independently integrated in the same *attB* site and despite 7 mismatches with the 14 bp *attB* from 8325-4, resolution to the original *attP* sequence occurred with the same precision as seen when Φ13kan^R^ excised from *attB* of 8325-4phi13kan^R^. In summary, our results demonstrate that excision of Φ13kan^R^ from alternative integration sites leads to evolutionary adaptation of the phage to the bacterium by increasing the number of *attP* nucleotides matching the alternative *attB* sequences.

### Phage adaptation to alternative *attB* site

After observing that induction of phages at alternative integration sites led to mutated phage populations with increased base pair matches between *attP* and the alternative *attB* sites or *attB_LA_*, we wondered whether these phages, in comparison to the original Φ13kan^R^, had increased preference for such sites in a new infection cycle. To address this we quantified integration by qPCR with primer pairs covering *attR*. We examined phage pools obtained from lysogen 2 and 7 (designated Φlys2 and Φlys7) excised from the *attB_LA_* and compared them to the original Φ13kan^R^ with respect to integration in either 8325-4 or 8325-4attBmut (Figure 3). As expected, we found that for the wildtype, homogeneous Φ13kan^R^ there were much less integration in *attB_LA_* compared to *attB* that matches the *attP* sequence. In contrast, this difference was essentially eliminated for the Φlys2 and Φlys7 phage pools. Further, the mutations in these pools significantly increased the integration frequency in 8325-4attBmut when compared to Φ13kan^R^ with the original *attP* site. Our results show that a single round of integration and excision dramatically increases the preference of the phage for an alternative or mutated attachment site.

**Figure 3.**
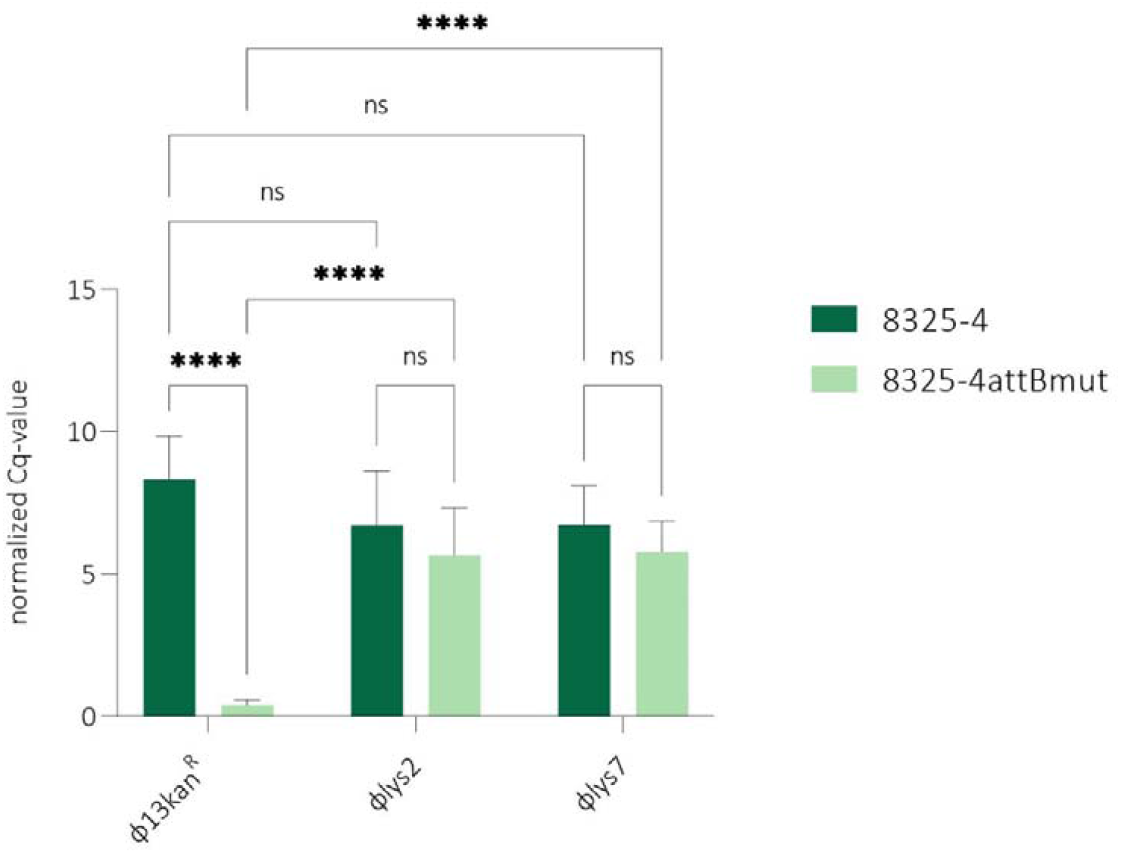
Normalized Cq-values after qPCR assay for detection of integration of evolved phages. The normalized Cq-value (calculated by 2^Cq(*pta*)-Cq(*hlb*)^) normalizes the cycle number of the gene of interest to the reference gene *pta*. Primers are identifying integration in *hlb (attB* or *attBA* for infection of 8325-4 (dark green) or 8325-4attBmut (light green) either with □ 13kan^R^ or the evolved phages □Tlys2 and □lys7, previously integrated at *attB_LA_*. Statistical analysis was carried out in GraphPad Prism 9.1.0, using Two-way ANOVA. *P* values: ns > 0,1234, *=0,0332, **=0,0021, ***=0,0002, ****<0,0001. Error bars represent standard deviation of three biological replicates with three technical replicates.

## Discussion

Sa3int prophages encode immune evasion factors and are found in most human strains of *S. aureus* ^18,19^. In contrast, LA-MRSA commonly lack Sa3int phages but when present, they increase the risk of transmission between household members and into the community ^3,7^. The integration site for Sa3int phages is naturally mutated in livestock-associated strains and so integration is infrequent and occurs at alternative sites often leading to mismatches between the *attL* and *attR* sequences. Intriguingly, we show that induction of these lysogens results in phage populations that are heterogeneous with respect to their *attP* sequences and with mutations that increase overall identity to the alternative bacterial integration site (Table 1,2 and Figure 1). Importantly, these *attP* changes increase phage integration into the naïve 8325-4attBmut in a new round of infection (Figure 3). Further we find that Sa3int prophages are spontaneously released from alternative integration sites highlighting that environmental triggers are not necessary for dissemination of these phages. Thus, rounds of excision and integration are possible with the potential for phage adaptation in each round.

When examining Sa3int prophages from outbreak strains of LA-MRSA ^7^ we observe a greater number of adaptive changes in the *attP* sites of the excised phages than from our model 8325- 4attBmut strain. This suggests that adapted phages are circulating in the LA-MRSA CC398 population, a notion that is supported by a study of the Sa3int phage P282 from a *S. aureus* CC398 strain, where the *attP* sequence is identical to *attB_LA_* ^15^. Also, re-analysis of Sa3Int-prophages in MRSA CC398 isolates from hospital patients in Germany ^16^ revealed that in 10 out of 15 lysogens, the *attL* and *attR* sequences were identical to *attB_LA_* (Supplementary Table S3) indicating that the prophages have adapted to the livestock-associated strains. This raises the question where phage adaptations may occur. Since about one in three humans is colonized with *S. aureus* of which the majority contains Sa3int phages, transfer can occur when humans, naturally colonized with *S. aureus*, are exposed to livestock-associated strains. Once established as a prophage in a livestock-associated strain, Sa3int phages will be released and, if adapted, will integrate more effectively than the original phage in the LA-MRSA population.

Integration at secondary sites has also been observed for other phages when the primary site is absent or mutated ^20–23^. Excision of phage λ from such as site resulted in substitutions in *attP* ^24,25^ and in P2, the authors stated that the new *attP* region contained DNA from *attR* ^26,27^. Similar to Φ13, these phages encode tyrosine recombinases ^12,22^. This family of recombinases catalyzes recombination between substrates with limited sequence identity ^28^. We propose that the adaptive behavior of Sa3int phages is depending on this promiscuity. As tyrosine-type recombinases are employed also by other *S. aureus* phages encoding virulence factors ^29^, the results presented here may provide a broader explanation for how phages adapt to new bacterial strains and thereby enable the host jumps that are regularly observed for staphylococci ^1^.

## Materials and Methods

### Strains and media

Phage-cured *S. aureus* 8325-4 ^30^ and its mutant 8325-4Φ13attBmut ^13^ (here termed 8325-4attBmut) containing the 2bp variation in *hlb* were used as recipients and indicator strains for Φ13kan^R^. Twenty LA-*S. aureus* strains harboring a Sa3Int-phages were analyzed for their *attR* and *attL* composition ^7^. Sequencing data available at https://www.ebi.ac.uk/ena/browser/home with identifiers listed in Supplementary Table S4. *S. aureus* S0385 (GenBank accession no. NC_017333) were used as a reference strain for analysis of sequencing data of the LA-strains. The prophage Φ13kan^R^ carries the kanamycin resistance cassette *aph*A3, which replaces the virulence genes *scn* and *chp* and was obtained by induction of 8325-4phi13kan^R^ ^13^. A full strain list is provided in Supplementary Table S1. Strains were grown in tryptone soy broth (TSB, CM0876, Oxoid) and tryptone soy agar (TSA, CM0131, Oxoid). Top agar for the overlay assays was 0,2 ml TSA/ml TSB. Kanamycin (30 μg/ml) and sheep blood agar (5%) were used to select for lysogens.

### Lysogenization assay

To obtain the phage stock, 8325-4phi13kan^R^ was grown to late exponential phase (37°C, 200 rpm, OD_600_=0,8), mixed with 2 μl/ml mitomycin C and incubated for another 2-4 hours. Phages were harvested by centrifugation for 5 min at 8150 x g and filtering the supernatant with a 0,2 μm membrane filter. The lysogens were obtained as described previously, with slight adjustments ^31^. In brief, Φ13kan^R^ was added at a multiplicity of infection MOI=1 to the respective recipients, incubated 30 min on ice to allow phage attachment, the non-attached phages were washed off and after another incubation for 30 min at 37°C allowing phage infection, the culture was diluted and plated on TSA with 5% blood and 30 μg/ml kanamycin. After overnight incubation at 37°C, 20 colonies showing β-hemolysis and two colonies without β-hemolysis were isolated and used for further analysis. Lysogens were derived from eight independent lysogenization experiments resulting in lysogens 1-5 (experiment 1); lysogens 6 and 7 (experiment 2), lysogen 8 (experiment 3), 10 and 11 (experiment 4), 12 and 13 (experiment 5), 14 and 15 (experiment 6), 16-19 (experiment 7) and lysogens 20-23 (experiment 8).

### Spot assay and phage propagation

Phage lysates were serially diluted in SM-buffer (100 mM NaCl, 50 mM Tris (pH=7,8), 1 mM MgSO_4_, 4 mM CaCl_2_) and spotted on recipient lawn of either *S. aureus* 8325-4 for pfu determination. To obtain an even lawn, 100 μl of fresh culture (OD=1) were added to 3 ml top agar and poured on a TSA-plate supplemented with 10mM CaCl_2_. After solidifying of the top agar, drops of 3 x 10 μl of each dilution were spotted on the lawn.

### Induction assay

To determine the different levels of phage release, the 8325-4attBmut-lysogens were grown to OD_600_=0,8, and centrifuged after adding 2 μg/ml mitomycin C and further incubation for 2 hours. The sterile-filtered supernatant was diluted and spotted on an overlay of 8325-4 consisting of 100 μl culture mixed with 3 ml top agar.

### Whole-genome sequencing and bioinformatics analysis

Genomic DNA was extracted by using DNeasy Blood and Tissue Kit (Qiagen) and whole genome sequences were obtained by 251bp paired-end sequencing (MiSeq, Illumina) as described previously ^32^. Raw data can be accessed at https://www.ebi.ac.uk/ena/browser/home with identifiers listed in Supplementary Table S5. Genomes were assembled using SPAdes ^33^. Geneious Prime 2020.1.1 was used to determine phage integration sites (www.geneious.com). The locations and core sequences were determined by extracting short sequences from the assembled draft genomes of the lysogens lying adjacent to the prophage and mapping it to the annotated genome of *S. aureus* 8325 (GenBank accession no. NC_007795). Reads obtained by sequencing the PCR amplicons spanning *attP*, were mapped to the Φ13 reference genome (GenBank accession no. NC_004617) and SNPs were called applying a variant frequency threshold of 50%. WebLogo3 was applied to detect gapped motifs in the flanking regions of the alternative *attB* sites ^34^.

### PCR and amplicon sequencing

Direct colony PCR was used to determine (i) the presence of the phage using *sak*-primers, (ii) the integrity of the *hlb* gene using *hlb*-primers and (iii) *attP* using *attP*st-primers ^35^ if the phage had spontaneously excised and was present in its circular form.

Primer sequences and cycling conditions are listed in Supplementary Table S2. For each reaction, a well-isolated colony was picked, suspended in 50 μl MilliQ-water, heat-lysed for 5 min at 99°C and briefly centrifuged. One μl was used as template. To determine *attP* of induced phages in lysates, 1 μl of a 1:10 dilution of phage lysate was used as template. Each single reaction mix was composed of 20,375 μl water, 2,5ml Taq polymerase buffer, 1 μl each of forward and reverse primers (10 μM), 0,5 μl dNTPs and 0,125 μl Taq polymerase (Thermo Fisher). PCR products were purified with GeneJET PCR purification kit (Thermo Fisher) and sequenced either using Sanger Sequencing (Mix2Seq, Eurofins Genomics) for the Sa3Int-phages deriving from the LA-MRSA strains or using Illumina MiSeq (sequencing depth varied from 10.000-180.000 (average 100.000)).

### qPCR assay

DNA for use in the qPCR assay (LightCycler 96, Roche) was extracted using the GenElute Bacterial Genomic DNA kit (Sigma). The samples of interest were obtained by lysogenizing *S. aureus* 8325-4 and 8325-4attBmut with the respective phage (Φ13kanR, Φlys2 or Φlys7) and plating 2 x 100 μl of the culture on TSA supplemented with 30μg/ml kanamycin. After overnight incubation, the colonies were scraped off (approx. 10.000 colonies) and re-suspended in 1 ml saline. Of this, 100 μl were used directly in the first lysis step of the kit. DNA concentration was measured using a Qubit^™^ (Invitrogen) and diluted to 1 ng/ml of which 5 μl were used in the qPCR reaction, consisting of 3 μl water, 10μl FastStart Essential DNA Green Master 2×, 1 μl of each forward and reverse primers (10 μM). Primer sequences and cycling conditions can be found in Supplementary Table S2.

## Author Contributions

H.L. and H.I. designed the study, H.L. generated experimental data, did formal analysis, wrote the manuscript and visualized the data; R.S. supported bioinformatic analysis; J.L. provided strain material; M.S. conducted sequencing; H.L., H.I., R.S., M.S. J.L. conducted review and editing; H.I. provided funding acquisition and project administration.

## Data availability

All genomic data used or produced in this study is deposited at the European Nucleotide Archive (https://www.ebi.ac.uk/ena/browser/home). Accession numbers and identifiers are listed in Supplementary Table S4 and S5.

Source data for the qPCR-assay and Sanger amplicon sequencing are provided with this paper.

## Acknowledgments

We thank Henrike Zschach for her contribution to the bioinformatic analysis and the staff of the Danish reference laboratory for Staphylococci at Statens Serum Institut for typing and handling of study isolates.

This project has received funding from the European Union’s Horizon 2020 research No. 765147

