## Supplementary information for "Bacteriophages encoding human immune evasion factors adapt to livestock-associated MRSA through rounds of integration and excision"

**Table S1** List of S. aureus strains used in this study.

| **Referred to in this study** | **Other name** | **Description** | **Reference** |
| --- | --- | --- | --- |
| 8325-4 |  | phage cured NCTC8325 | 30 |
| 8325-4attBmut | 8325-4ɸ13attBmut | 8325-4 mutated at ɸ13 attB site in hlb, hlb+ | 13 |
| 8325-4ɸ13kan^R^ |  | 8325-4 lysogenized with ɸ13kanR, hlb-, kanamycin resistant | 13 |
| Variant I | 123193 | CC398, Sa3int phage intergenic between SAPIG_RS03725 and SAPIG_RS03730 | 7 |
| Variant I | 146281 | CC398, Sa3int phage intergenic between SAPIG_RS03725 and SAPIG_RS03730 | 7 |
| Variant I | 147327 | CC398, Sa3int phage intergenic between SAPIG_RS03725 and SAPIG_RS03730 | 7 |
| Variant I | 147333 | CC398, Sa3int phage intergenic between SAPIG_RS03725 and SAPIG_RS03730 | 7 |
| Variant I | 147335 | CC398, Sa3int phage intergenic between SAPIG_RS03725 and SAPIG_RS03730 | 7 |
| Variant II | 147331 | CC398, Sa3int phage in sfaC SAPIG_RS11745 | 7 |
| Variant II | 147339 | CC398, Sa3int phage in sfaC SAPIG_RS11745 | 7 |
| Variant III | 157445 | CC398, Sa3int phage in sph SAPIG_RS10795 | 7 |
| Variant III | 157659 | CC398, Sa3int phage in sph SAPIG_RS10795 | 7 |
| Variant III | 157661 | CC398, Sa3int phage in sph SAPIG_RS10795 | 7 |
| Variant III | 157663 | CC398, Sa3int phage in sph SAPIG_RS10795 | 7 |
| Variant IV | 155519 | CC398, Sa3int phage in cidA SAPIG_RS13630 | 7 |
| Variant V | 153891 | CC398, Sa3int phage in sph SAPIG_RS10795 | 7 |
| Variant VI | 154501 | CC398, Sa3int phage in CocE/NonD SAPIG_RS13905 | 7 |
| Variant VI | 154789 | CC398, Sa3int phage in CocE/NonD SAPIG_RS13905 | 7 |
| Variant VI | 154791 | CC398, Sa3int phage in CocE/NonD SAPIG_RS13905 | 7 |
| Variant VI | 154923 | CC398, Sa3int phage in CocE/NonD SAPIG_RS13905 | 7 |
| Variant VI | 55-103-045 | CC398, Sa3int phage in CocE/NonD SAPIG_RS13905 | 7 |
| Variant VI | 55-103-046 | CC398, Sa3int phage in CocE/NonD SAPIG_RS13905 | 7 |
| Variant VI | 55-103-047 | CC398, Sa3int phage in CocE/NonD SAPIG_RS13905 | 7 |
| Lysogen 1 |  | 8325-4attBmut with ɸ13kan^R^ in SAOUHSC_01067 CDS conserved hypothetical protein | this study |
| Lysogen 2 |  | 8325-4attBmut with ɸ13kan^R^ in hlb sphingomyelin phosphodiesterase | this study |
| Lysogen 3 |  | 8325-4attBmut with ɸ13kan^R^ in SAOUHSC_02680 narY nitrate reductase subunit beta | this study |
| Lysogen 4 |  | 8325-4attBmut with ɸ13kan^R^ inSAOUHSC_01263 phosphodiesterase | this study |
| Lysogen 5 |  | 8325-4attBmut with ɸ13kan^R^ intergenic between hypothetical proteins SAOUHSC_01301 and SAOUHSC_01304 | this study |
| Lysogen 6 |  | 8325-4attBmut with ɸ13kan^R^ in SAOUHSC_02396 hypothetical protein | this study |
| Lysogen 7 |  | 8325-4attBmut with ɸ13kan^R^ in hlb sphingomyelin phosphodiesterase | this study |
| Lysogen 8 |  | 8325-4attBmut with ɸ13kan^R^ in SAOUHSC_00952 hypothetical protein | this study |
| Lysogen 10 |  | 8325-4attBmut with ɸ13kan^R^ in SAOUHSC_00125 cap5L protein/glycosyltransferase | this study |
| Lysogen 11 |  | 8325-4attBmut with ɸ13kan^R^ in SAOUHSC_00436 gltD glutamate synthase subunit beta | this study |
| Lysogen 12 |  | 8325-4attBmut with ɸ13kan^R^ in SAOUHSC_02433 hypothetical protein | this study |
| Lysogen 13 |  | 8325-4attBmut with ɸ13kan^R^ intergenic between hypothetical proteins SAOUHSC_01301 and SAOUHSC_01304 | this study |
| Lysogen 14 |  | 8325-4attBmut with ɸ13kan^R^ in SAOUHSC_01067 CDS conserved hypothetical protein | this study |
| Lysogen 15 |  | 8325-4attBmut with ɸ13kan^R^ in SAOUHSC_02428 hypothetical protein | this study |
| Lysogen 16 |  | 8325-4attBmut with ɸ13kan^R^ in SAOUHSC_02436 hypothetical protein | this study |
| Lysogen 17 |  | 8325-4attBmut with ɸ13kan^R^ in SAOUHSC_02430 ABC transporter substrate-binding protein CDS | this study |
| Lysogen 18 |  | 8325-4attBmut with ɸ13kan^R^ in SAOUHSC_01067 CDS conserved hypothetical protein | this study |
| Lysogen 19 |  | 8325-4attBmut with ɸ13kan^R^ in SAOUHSC_00910 hypothetical protein CDS | this study |
| Lysogen 20 |  | 8325-4attBmut with ɸ13kan^R^ in SAOUHSC_02990 hypothetical protein CDS | this study |
| Lysogen 21 |  | 8325-4attBmut with ɸ13kan^R^ in SAOUHSC_00125 cap5L protein/glycosyltransferase | this study |
| Lysogen 22 |  | 8325-4attBmut with ɸ13kan^R^ in SAOUHSC_00395 superantigen-like protein CDS | this study |
| Lysogen 23 |  | 8325-4attBmut with ɸ13kan^R^ in SAOUHSC_00910 hypothetical protein CDS | this study |

**Table S2** Overview of used primers and cycling conditions for conventional and qPCR. If not otherwise stated, the primers have been designed for this study.

| **Primer name** | **Sequence 5´- 3´** | **Annealing temp. °C** | **Elongation time (s)** | **Reference** |
| --- | --- | --- | --- | --- |
| hlb_fwd | ATGGTGAAAAAAACAAAATCCAATTCAC | 50 | 60 |  |
| hlb_rev | CTATTTACTATAGGCTTTGATTGGGTAATG |  |  |  |
| Sak_fwd | GTGCATCAAGTTCATTCGAC | 49 | 30 | Tang et al., 2017 |
| Sak_rev | TAAGTTGAATCCAGGGTTTT |  |  |  |
| attPst_fwd | TCTAGCTTTTGGGGTGTACATTCC | 49 | 30 | Goerke et al., 2006 |
| attPst_rev | GCTTTGAAATCAGCCTGTAGAG |  |  |  |
| Pta fwd | AGAAGCAATCATTGATGGCGA | 55 | 10 | Aedo and Tomasz, 2016 |
| Pta rev | ACCTGGCGCTTTTTTCTCAG |  |  |  |
| Φ13 attR fwd | CTCCAAACCCAATAAATACTGTTGTTAC | 55 | 10 |  |
| Hlb attR rev | CGAGTACAGGTGTTTGATAAGGATATTC | 55 | 10 |  |

**Table S3** Alternative integration sites extracted from previous studies. In the study by Kraushaar et al., the authors stated attB sequences that refered to attR, hence the information is given in grey font but kept to show the alternative sites. For the study by Goerke et al., no sequence data was available.

| **Alternative *attB*** | **Location** | **Reference genome** | **Accession no.** | **Ref.** |
| --- | --- | --- | --- | --- |
| TGTATCCAAACTGG | Original *attB* in *hlb* of *S. aureus* 8325-4 |  |  |  |
| **GT**TATCCAA**T**CTGG | alanine racemase | *S. aureus* CC398, 61599 | PRJEB25608 | 13 |
| **G**G**GGA**CC**T**AACTGG | nitrate reductase |  |  |  |
| **CCAT**TCCA**T**ACTGG | ornithine carbamoyltrasferase |  |  |  |
| **GTGTAT**CCA**T**CTGG | FADH(2)-oxidizing methlentetrahydrofolate-tRNA-(uracil(54)-C(5))-methyltransferase TrmFO) |  |  |  |
| T**T**TATC**GTTT**CTGG | Acycl ersterase |  |  |  |
| T**T**TATCC**GT**A**A**TG**C** | between SAPIG2164: aldehyde dehydrogenase family protein and SPIG2165: HxlR family transcriptional regulator |  |  |  |
| TGT**TCTTT**A**T**CTGG | plasmid JQ861959 integrated upstream of the *muTB* gene |  |  |  |
| **GT**T**TCT**C**C**A**C**CTGG | membrane protein (2x) |  |  |  |
| **C**G**ATG**CCAAA**A**TGG | pyruvate oxigenase | *S. aureus* ST398 | AM990992.1 | 16 |
| TGTA**C**C**TT**A**TGA**GG | *ilvB* |  |  |  |
| **CTAG**TCC**TT**ACTGT | between SAPIG0723 and SAPIG0724 (2x, different *attR* and *attL*, indicating different *attP*) |  |  |  |
| **GT**TATCCAA**T**CTGG | alanine racemase |  |  |  |
| TGTATCC**G**AA**T**TGG | *hlb* (10 isolates where *attR* and *attL* are identical to *attB_LA_*, 5 isolates where *attL=attB_LA_* and attR=attB, indicating different *attP*) |  |  |  |
| TG**AT**TCCAA**C**C**G**GG | Hypothetical protein StauST398-5_0050 | phage StauST398-5 | KC595279 |  |
| TGTATCC**G**AA**T**TGG | *hlb* |  | CP019593 | 14 |
| **CC**T**T**TCCA**T**A**A**TGG | Alpha/beta hydrolase |  | CP040229 |  |
| **AC**T**TC**CCAAACTGG | cytochrome d ubiquinol oxidase subunit II |  | CP040232 |  |
| TGTATCCAAACTGG | General stress protein (AUC50_04635) | *S. aureus* RIVM3897 | CP013621 | 15 |
| TGTATCCAA**T**CTGG | General stress protein (AUC50_04635) |  |  |  |
| TGTATCC**TT**ACTG**T** | Nucleoside permease (AUC50_03495) |  |  |  |
| TGTATCCAAACTGG | Cytochrome D ubiquinol oxidase subunit I (AUC50_05470) |  |  |  |
| TGTATCCAAACTG**A** | Hypothetical protein (AUC50_10740) |  |  |  |
| TGTATCCAAACTG**A** | GMP synthetase (AUC50_02200) |  |  |  |
| TGTATCCAA**G**CTGG | Gamma-aminobutyrate permease (AUC50_08995) (3x) |  |  |  |
| TGTATCCAAACTGG | Hypothetical protein (SAMI_1999) | *S. aureus* MI | AP017320 |  |
| TGTATCCA**T**ACTGG | Alanine racemase (A7327_11665) | *S. aureus* 08–02300 | CP015646 |  |
| TGTATCCAAACTGG | Integrase | *S. aureus* Sa54 | KT253891 |  |
|  | ORF SA1270 | *S. aureus* N315 | BA000018.3 | 35 |
| n. a. | intergenic between ORF SA0966 and SA0967 |  |  |  |
|  | *capL* |  |  |  |

**Table S4** Accession numbers and identifiers for source data of Table 1. Bioproject/biosample accessible at https://www.ebi.ac.uk/ena/browser/home. Mix2Seq Sanger Sequencing identifiers for PCR fragments containing attP of the induced Sa3Int-phages, data available with this paper.

| **Name used in this study** | **Study ID** | **Other ID** | **Bioproject** | **Biosample** | **Mix2Seq number** |
| --- | --- | --- | --- | --- | --- |
| Variant I | 123193 | SSI_123193 | PRJEB25608 | ERR1992266 | EF30859333 |
| Variant I | 146281 | SSI_146281 | PRJNA613886 | SRR11364549 | EF30859334 |
| Variant I | 147327 | SSI_147327 | PRJNA613886 | SRR11364546 | EF30859336 |
| Variant I | 147333 | SSI_147333 | PRJNA613886 | SRR11364544 | EF30859338 |
| Variant I | 147335 | SSI_147335 | PRJNA613886 | SRR11364543 | EF31211630 |
| Variant II | 147331 | SSI_147331 | PRJNA613886 | SRR11364545 | EF30859337 |
| Variant II | 147339 | SSI_147339 | PRJNA613886 | SRR11364542 | EF31211645 |
| Variant III | 157445 | SSI_157445 | PRJNA613886 | SRR11364487 | EF31634870 |
| Variant III | 157659 | SSI_157659 | PRJNA613886 | SRR11364554 | EF31634871 |
| Variant III | 157661 | SSI_157661 | PRJNA613886 | SRR11364553 | EF31634872 |
| Variant III | 157663 | SSI_157663 | PRJNA613886 | SRR11364552 | EF31634874 |
| Variant IV | 155519 | SSI_155519 | PRJNA613886 | SRR11364507 | EF31634869 |
| Variant V | 153891 | SSI_153891 | PRJNA613886 | SRR11364516 | EF31211647 |
| Variant VI | 154501 | SSI_154501 | PRJNA613886 | SRR11364513 | EF31634868 |
| Variant VI | 154789 | SSI_154789 | PRJNA613886 | SRR11364512 | EF31211648 |
| Variant VI | 154791 | SSI_154791 | PRJNA613886 | SRR11364511 | EF31203716 |
| Variant VI | 154923 | SSI_154923 | PRJNA613886 | SRR11364510 | EF31888721 |
| Variant VI | 55-103-045 |  | PRJEB25608 | ERR1992132 | EF31211652 |
| Variant VI | 55-103-046 |  | PRJEB25608 | ERR1992133 | EF31211815 |
| Variant VI | 55-103-047 |  | PRJEB25608 | ERR1992134 | EF31211656 |

**Table S5** Accession numbers and identifiers for the genomic data generated in this study. Raw reads available at https://www.ebi.ac.uk/ena/browser/home with Bioproject number PRJEB44479.

| **Name used in this study** | **Biosample** |
| --- | --- |
| Lysogen 1 | SAMEA8606235 |
| Lysogen 2 | SAMEA8606236 |
| Lysogen 3 | SAMEA8606237 |
| Lysogen 4 | SAMEA8606238 |
| Lysogen 5 | SAMEA8606239 |
| Lysogen 6 | SAMEA8606240 |
| Lysogen 7 | SAMEA8606229 |
| Lysogen 8 | SAMEA8606230 |
| Lysogen 10 | SAMEA8606231 |
| Lysogen 11 | SAMEA8606232 |
| Lysogen 12 | SAMEA8606233 |
| Lysogen 13 | SAMEA8606234 |
| Lysogen 14 | SAMEA8606241 |
| Lysogen 15 | SAMEA8606242 |
| Lysogen 16 | SAMEA8606243 |
| Lysogen 17 | SAMEA8606244 |
| Lysogen 18 | SAMEA8606245 |
| Lysogen 19 | SAMEA8606246 |
| Lysogen 20 | SAMEA8606247 |
| Lysogen 21 | SAMEA8606248 |
| Lysogen 22 | SAMEA8606249 |
| Lysogen 23 | SAMEA8606250 |
| attP Φlys1 | SAMEA8606266 |
| attP Φlys2 | SAMEA8606267 |
| attP Φlys3 | SAMEA8606251 |
| attP Φlys4 | SAMEA8606268 |
| attP Φlys5 | SAMEA8606253 |
| attP Φlys6 | SAMEA8606255 |
| attP Φlys7 | SAMEA8606257 |
| attP Φlys8 | SAMEA8606259 |
| attP Φlys10 | SAMEA8606269 |
| attP Φlys11 | SAMEA8606261 |
| attP Φlys12 | SAMEA8606270 |
| attP Φlys13 | SAMEA8606263 |
| attP Φlys14 | SAMEA8606265 |
| attP Φlys15 | SAMEA8606271 |
| attP Φlys16 | SAMEA8606252 |
| attP Φlys17 | SAMEA8606254 |
| attP Φlys18 | SAMEA8606256 |
| attP Φlys19 | SAMEA8606258 |
| attP Φlys20 | SAMEA8606260 |
| attP Φlys21 | SAMEA8606262 |
| attP Φlys22 | SAMEA8606264 |
| attP Φlys23 | SAMEA8606272 |
| attP Φphi13kan^R^ | SAMEA8606273 |

1	2	3	4	5	6	7	8	10	11	12	13	14 15	x	16 17 18 19	20	x	21	x	22 23

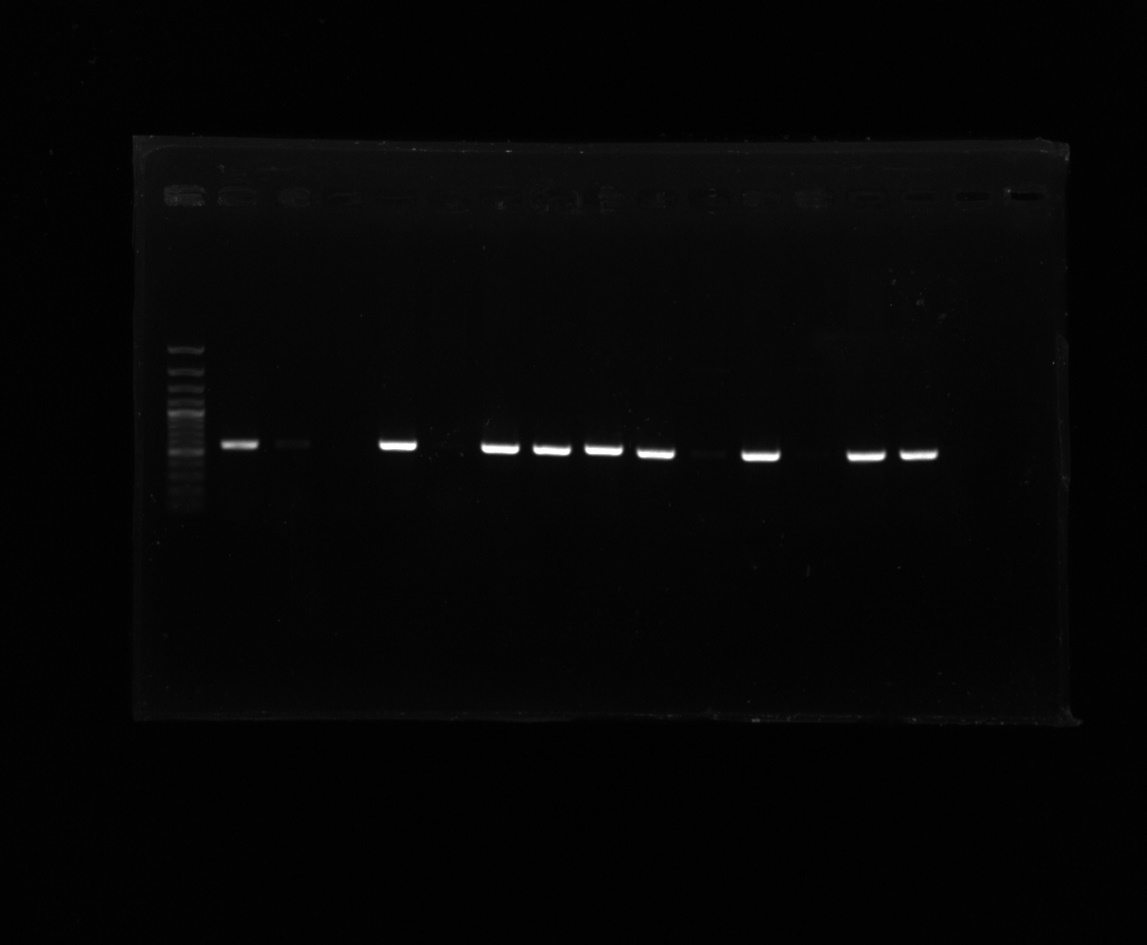

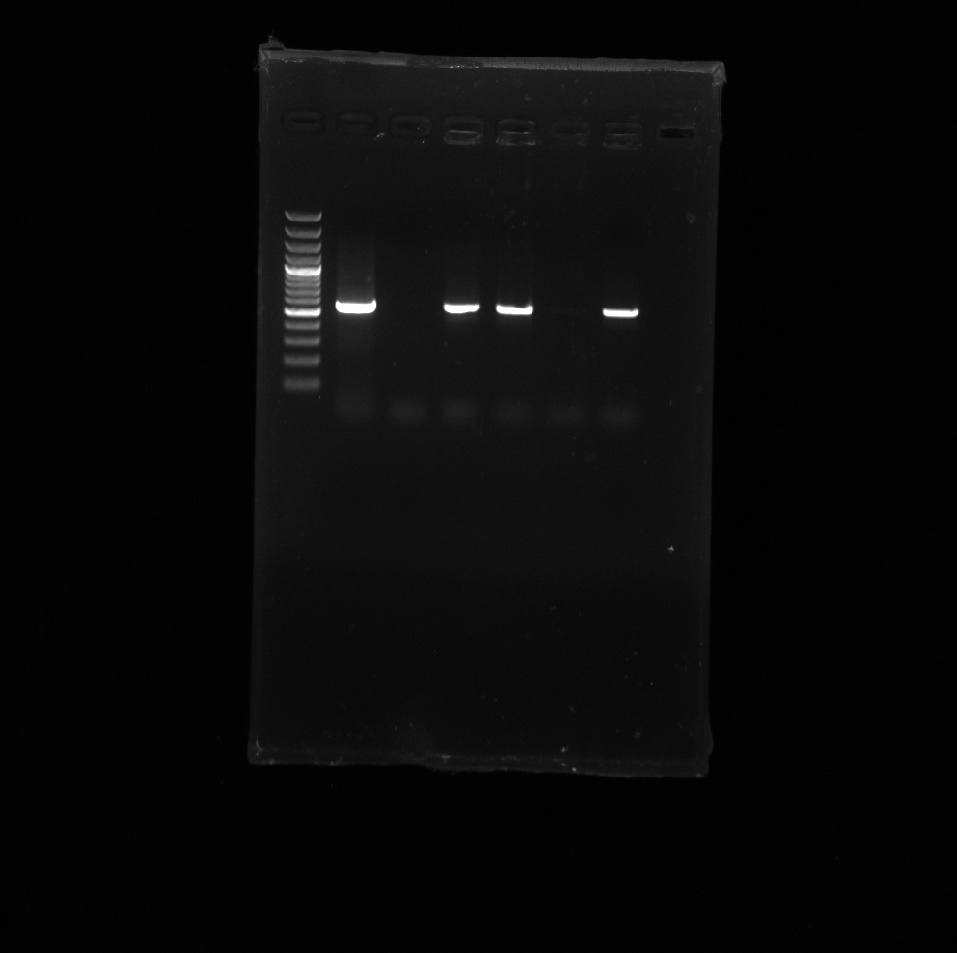

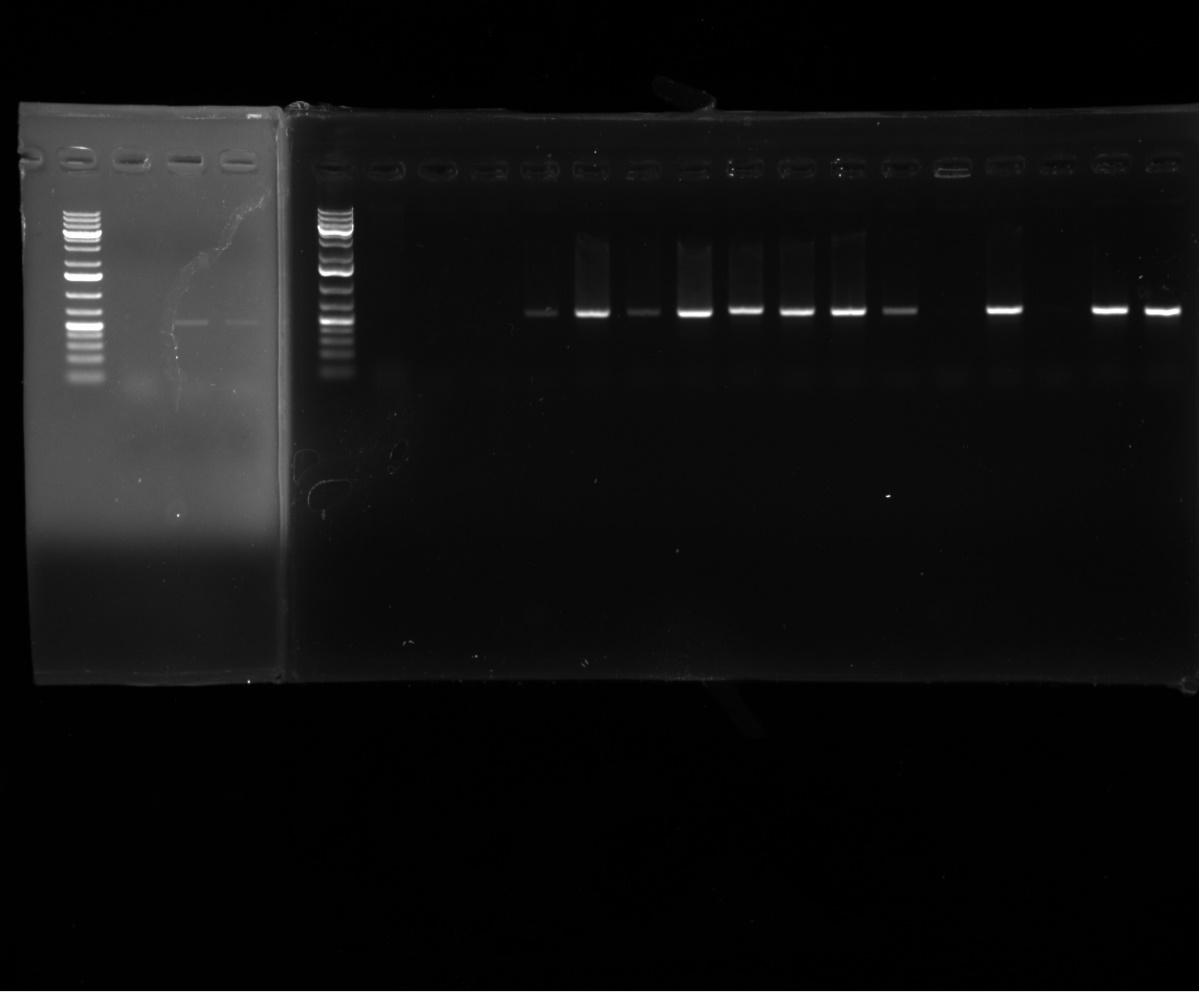

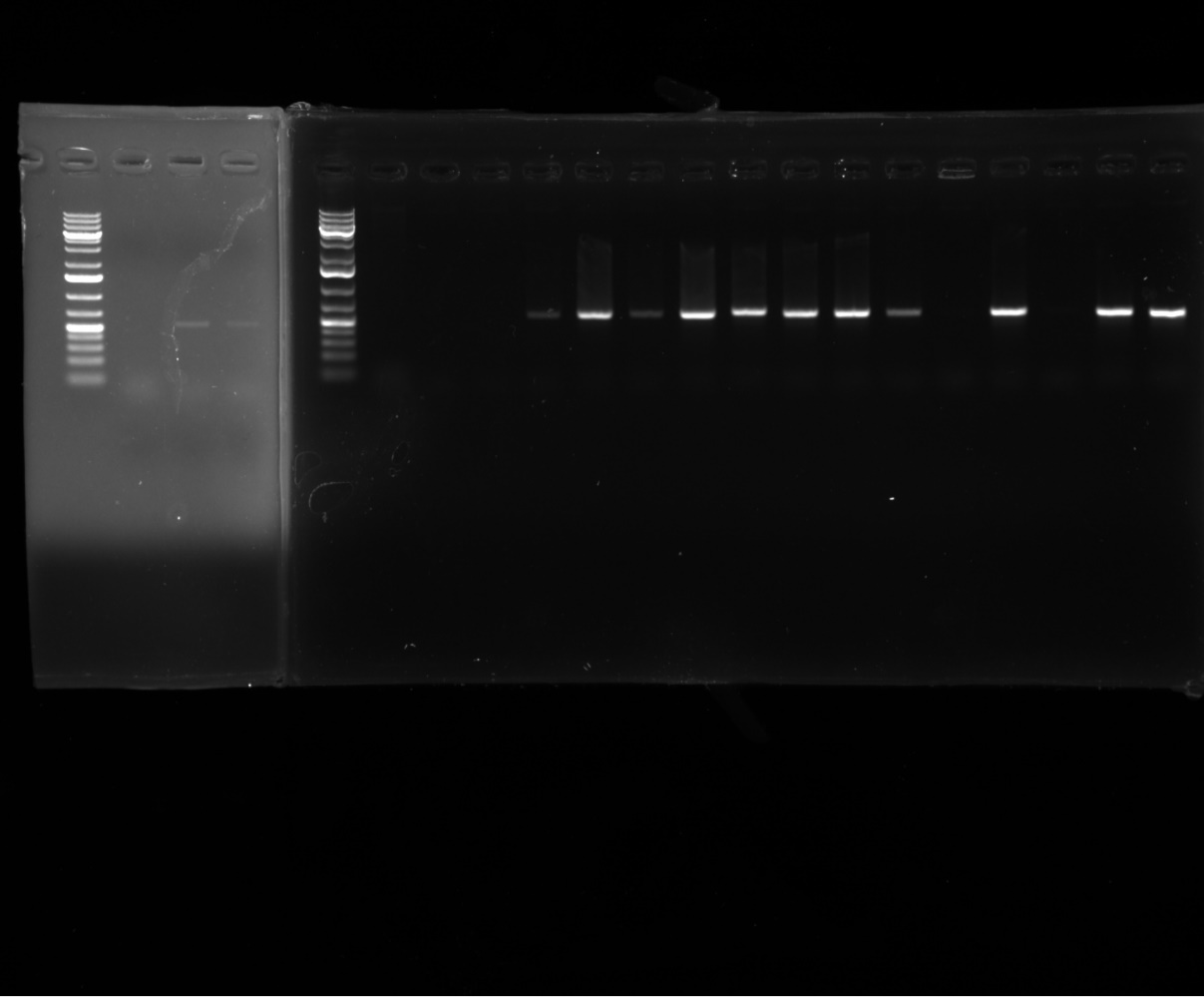

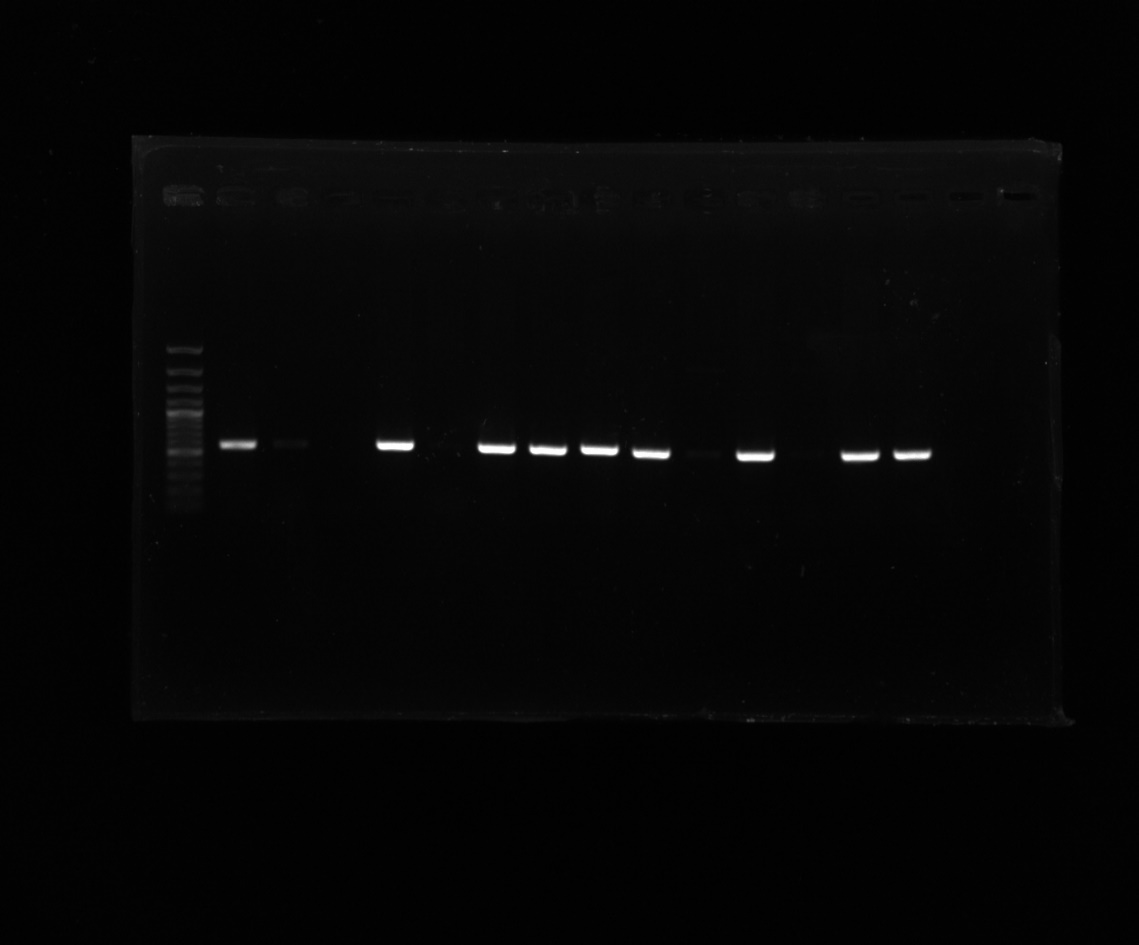

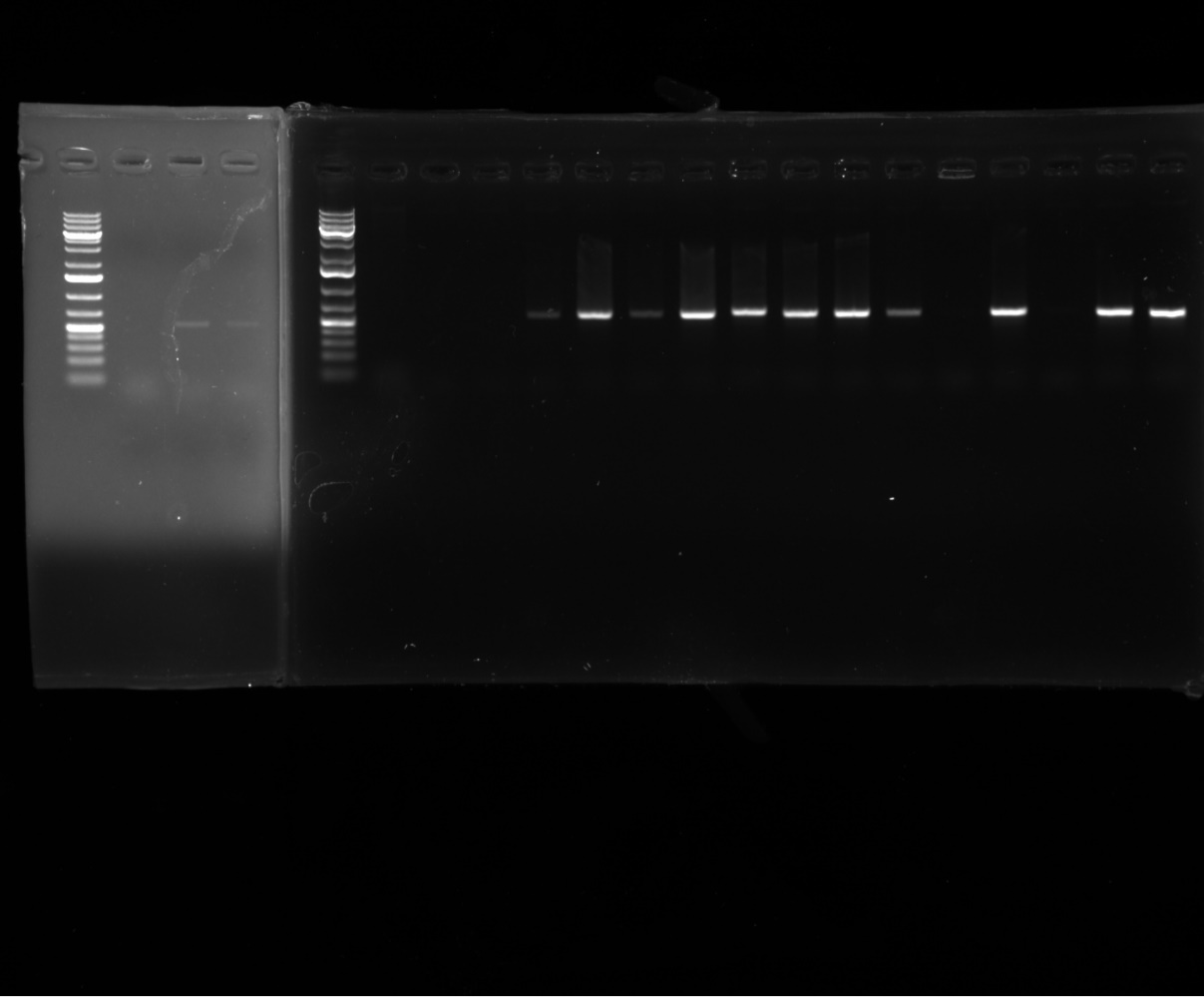

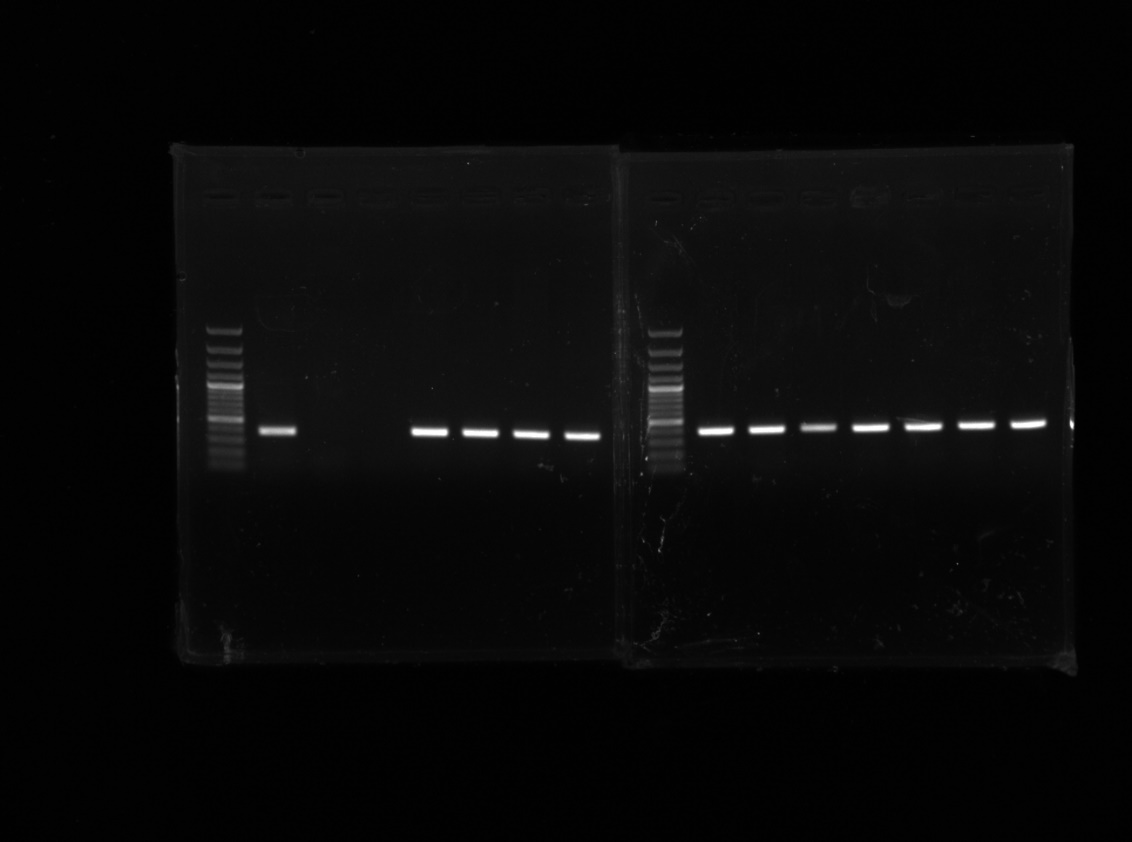

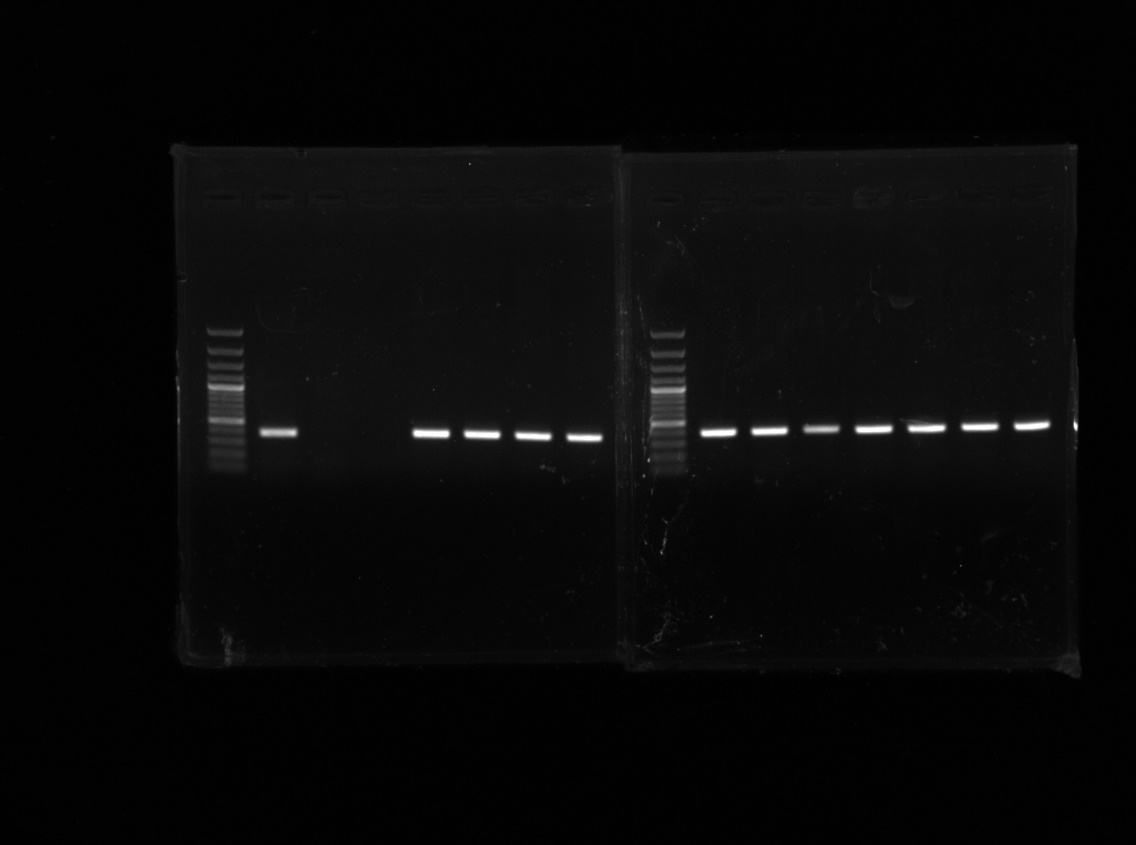

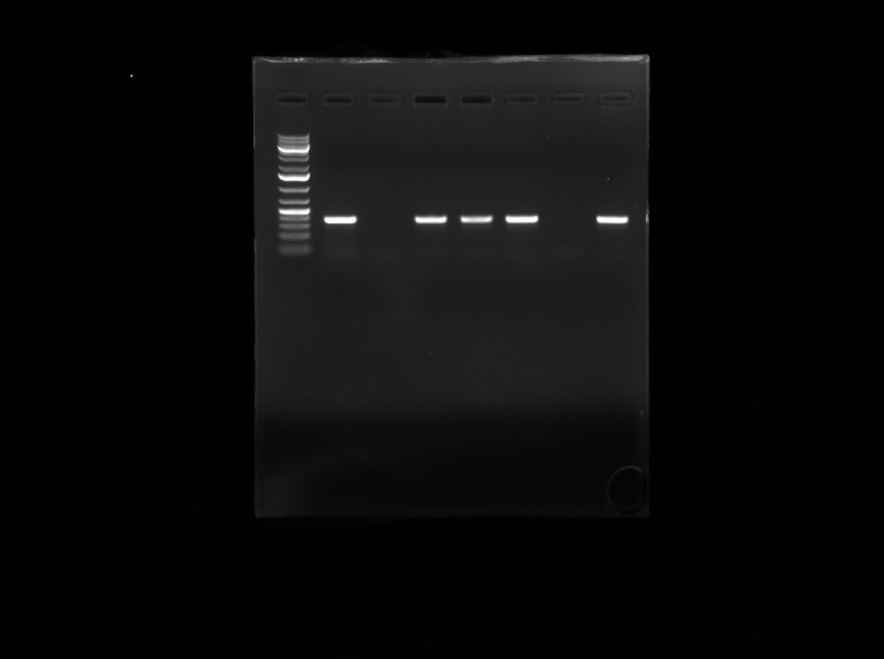

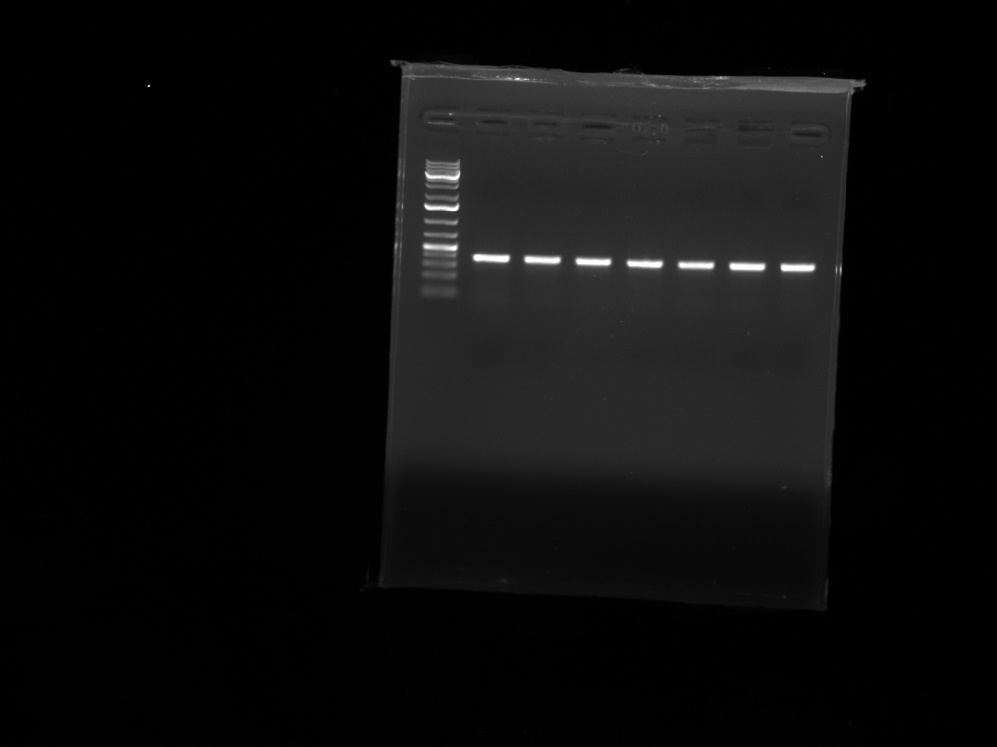

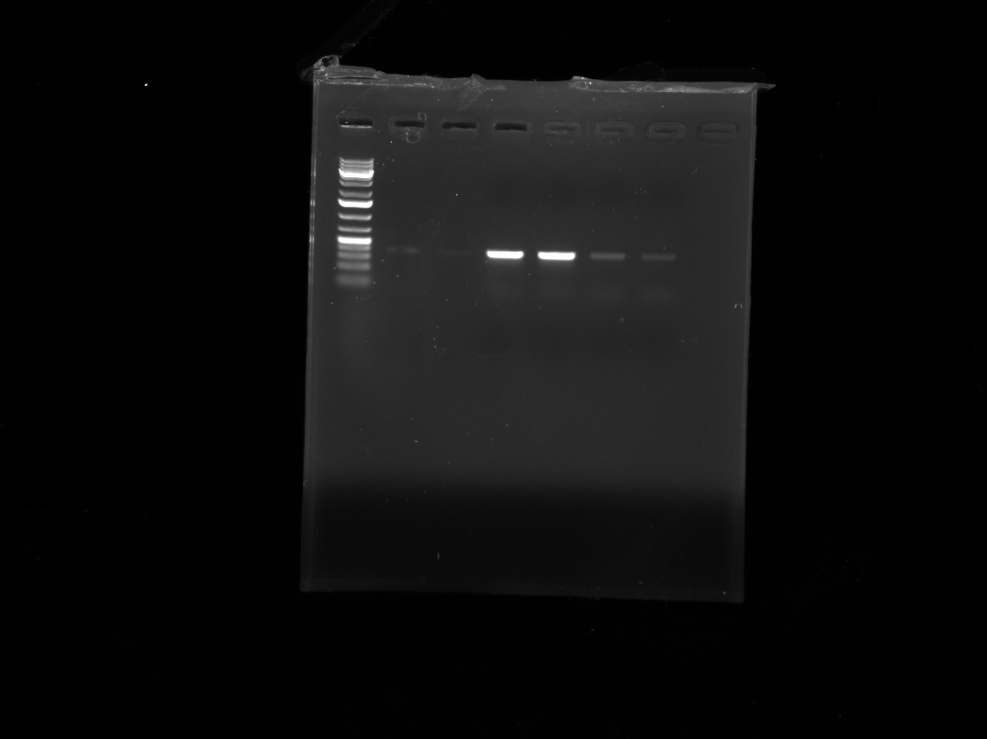

1	2	3	4	5	6	7	8	10 11	14 15	x	16	17 18	19 20	x	21	22 23

**Figure S1** Gelelectrophoresis after PCR for hlb (top) and sak (bottom). The pictures have been taken from different rounds of PCR amplification (as indicated by the white cuts between the pictures) and therefor are not completely aligned. Ladder is 1kb+ GeneRuler from Thermo Fisher Scientific. Lane numbers indicate lysogen number. Lanes marked with x are either empty or contain an unrelated sample.

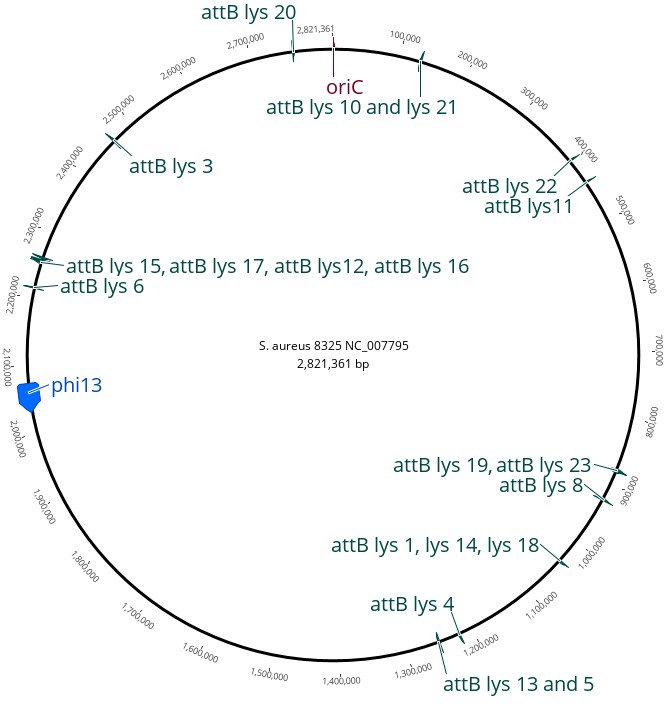

**Figure S2** Location of the alternative integration sites of Φ13 indicated in the reference chromosome of S. aureus 8325 (NC_007795.1). Figure generated using Geneious Prime 2021.1.1.

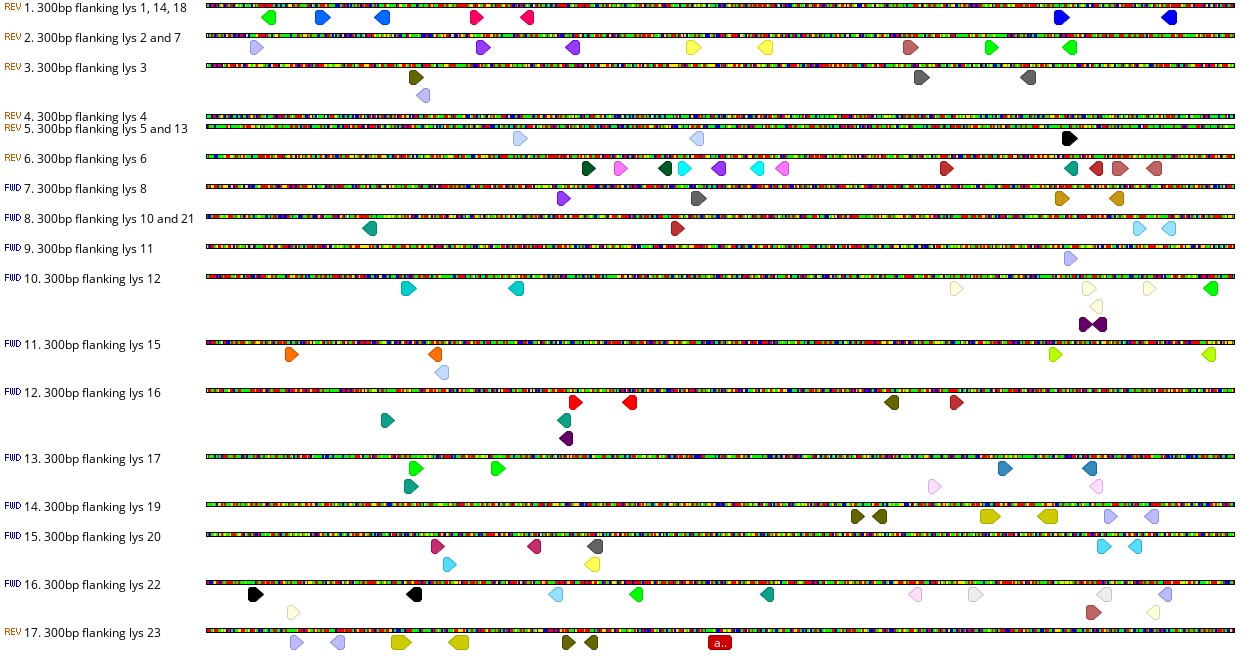

**Figure S3** Inverted and direct repeats of >8bp found in the 300bp flanking sequence of the 17 alternative attB sites in S. aureus 8325-4attBmut. Each identical sequence is highlighted in the same color. The red box in the center marks the 14bp attB sequence. The search has been carried out with repeat finder in Geneious Prime 2021.1.1

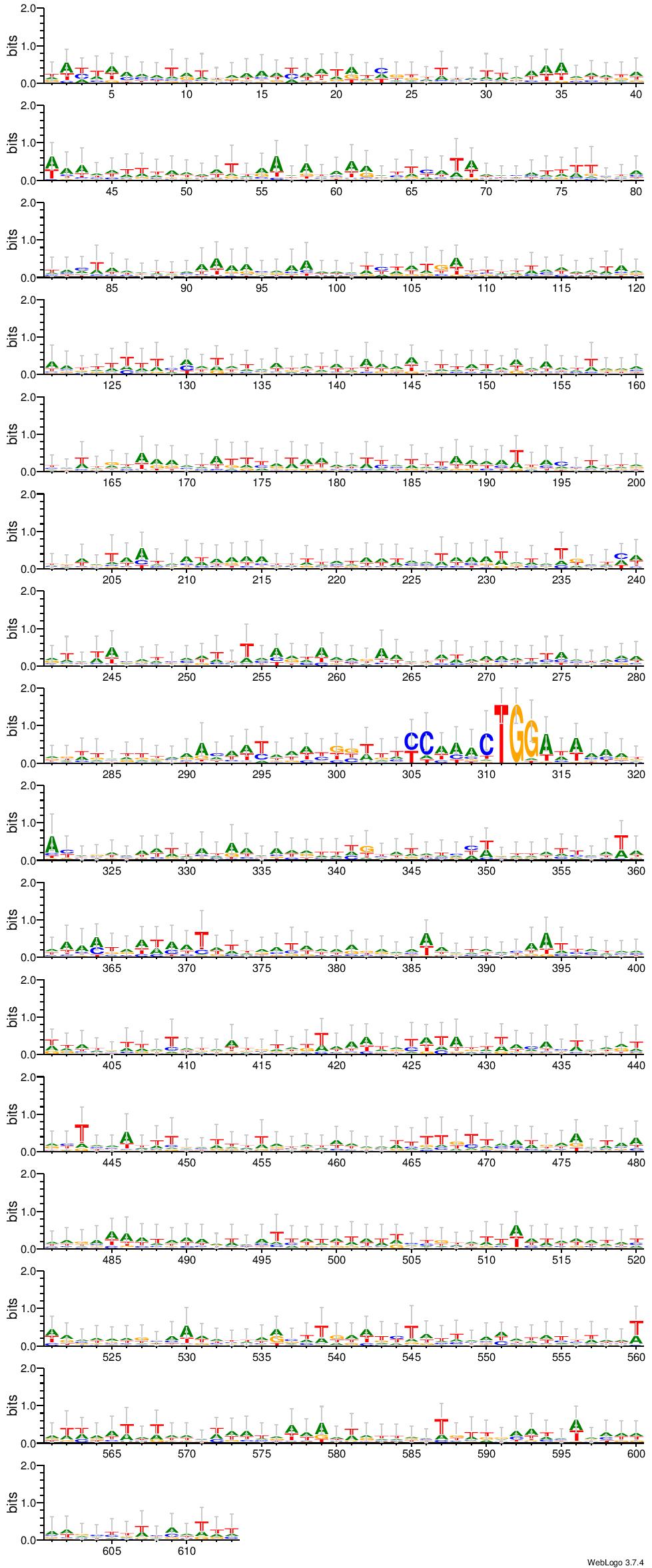

**Figure S4** Sequence logo of a sequence alignment of 300bp flanking the 17 alternative attB sites of S. aureus 8325-4attBmut. The 14bp attB site is highlighted with a square box. Each stack of letters represents one position in the DNA sequence. The height of the stack indicates the sequence conservation at the respective position, and the size of the letter within the stack represents the relative frequency of each nucleotide at this position. The logo was created using weblogo.threeplusone.com.

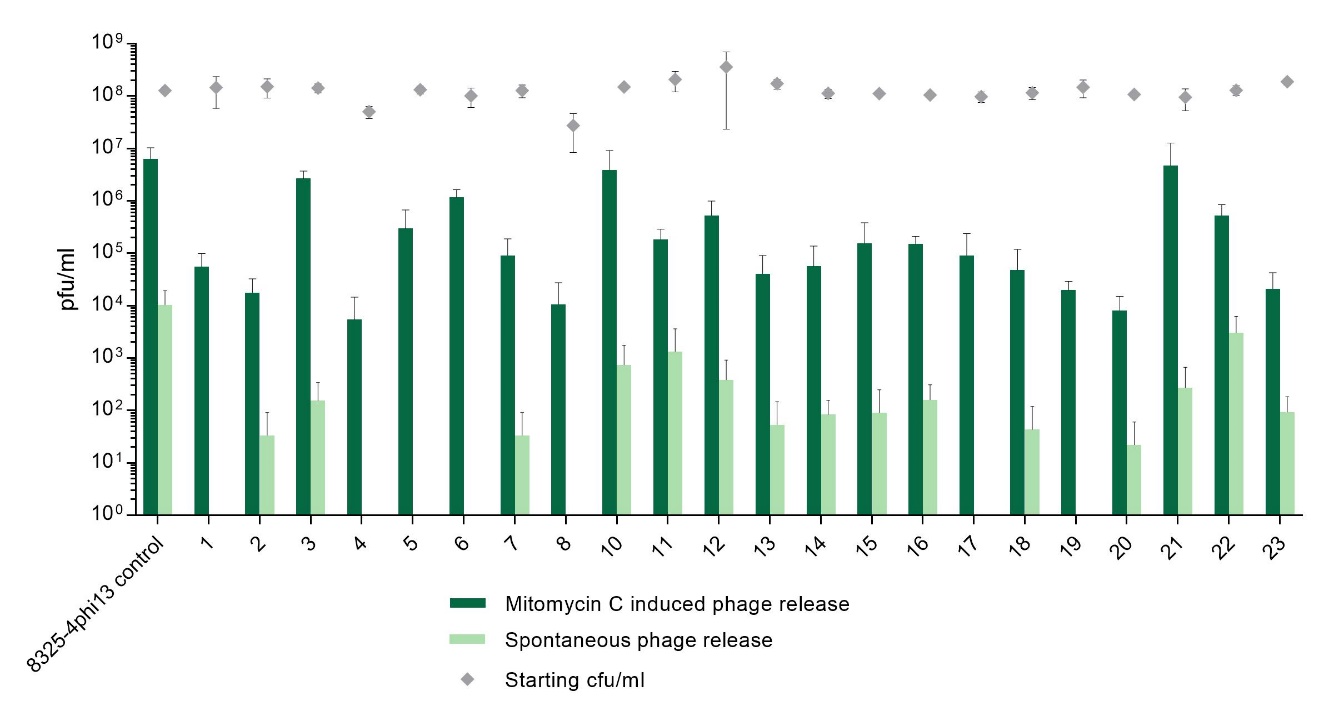

**Figure S5** Excision of Φ13kan^R^ from alternative integration sites. Plaque forming units (pfu/ml) observed from the 22 lysogens after spontaneous release (light green bars) or when induced with 2 µg/ml mitomycin C (dark green bars). The starting cell count (cfu/ml) is shown (grey diamonds) to exclude differences in pfu due to varying inoculum. Error bars represent standard deviation of three biological repeats.

**a**

**bA**

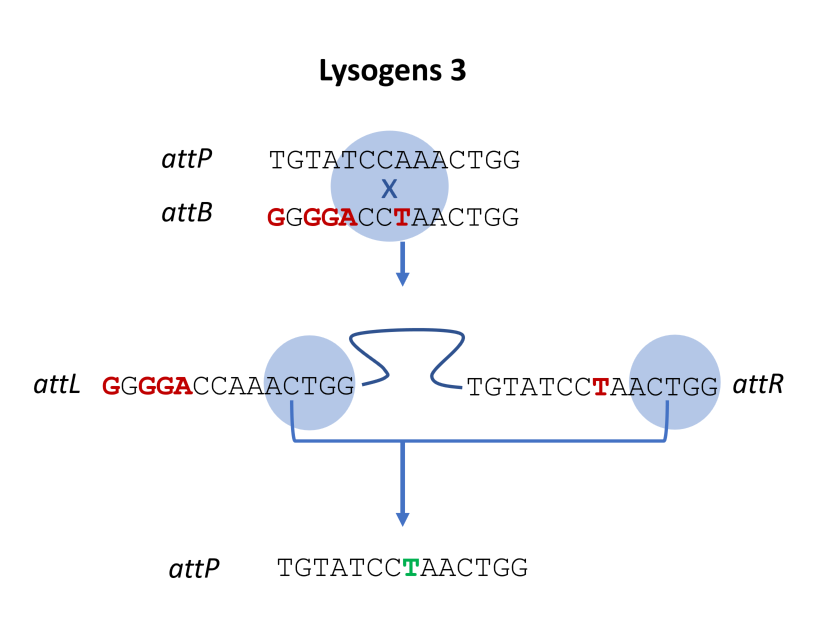

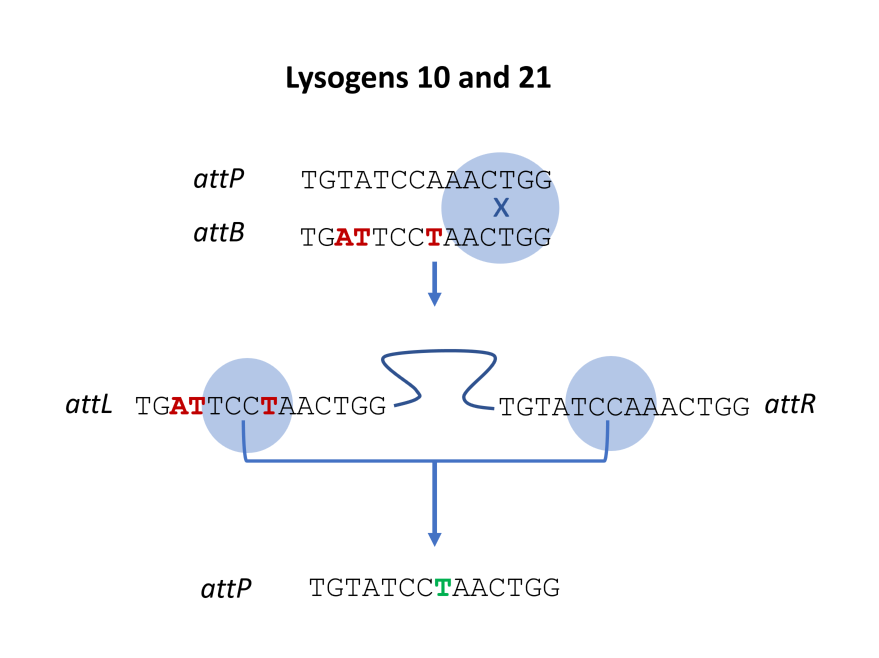

**Figure S6a** displays an integration event at the 3’-end of the att-sites and excision happening towards the 5' end (example of lysogen 10, 21). In **Figure S6b** displays integration at the center of att sites and excision at the 3' end (example of lysogen 3). Blue bubble indicate where the recombination events are taking place.
